# Development of an inhibitory TTC7B selective nanobody that blocks EFR3 recruitment of PI4KA

**DOI:** 10.1101/2025.07.28.667261

**Authors:** Sushant Suresh, Alexandria L Shaw, Damilola K Akintola, Martine Lunke, Sophia Doerr, Pooja Rohilla, Tamas Balla, Calvin K Yip, Scott D Hansen, Jennifer A Cobb, John E Burke

## Abstract

Phosphatidylinositol 4 kinase IIIα (PI4KIIIα/PI4KA) is an essential lipid kinase that plays a critical role in regulating plasma membrane identity. PI4KA is primarily recruited to the plasma membrane through the targeted recruitment by the proteins, EFR3A and EFR3B, which bind to the PI4KA accessory proteins TTC7 (TTC7A/B) and FAM126 (FAM126A/B). Here we characterised how both EFR3 isoforms interact with all possible TTC7-FAM126 combinations and developed a nanobody that specifically blocked EFR3-mediated PI4KA recruitment in TTC7B containing complexes. Most EFR3-TTC7-FAM126 combinations show similar binding affinities, with the exception of EFR3A-TTC7B-FAM126A, which binds with a ∼10-fold higher affinity. Moreover, we showed that EFR3B phosphorylation markedly decreased binding to TTC7-FAM126. Using a yeast display approach, we isolated a TTC7B selective nanobody that blocked EFR3 binding. Cryo-electron microscopy and hydrogen deuterium exchange mass spectrometry showed an extended interface with both PI4KA and TTC7B that sterically blocks EFR3 binding. The nanobody caused decreased membrane recruitment both on lipid bilayers and in cells, with decreased PM production of PI4P. Collectively, these findings provide new insights into PI4KA regulation and provide a tool for manipulating PI4KA complexes, that may be valuable for therapeutic targeting.

## Introduction

Phosphoinositides are critical mediators of many biological processes at the plasma membrane (PM), with the lipid phosphatidylinositol 4-phosphate (PI4P) being an essential PM lipid that is generated by the lipid kinase type III phosphatidylinositol 4 kinase (referred to by its gene name PI4KA) (1–3). PI4P is essential for defining the lipid identity of the PM (4), and it is the precursor for the downstream signaling lipids, PIP_2_ and PIP_3_, which play crucial roles in multiple signaling processes (5). The generation of PI4P at the PM is essential for enrichment of phosphatidylserine (PS) as PI4P provides the energy necessary for transport of PS from the ER against its concentration gradient (6). The predominant form of PI4KA exists as a complex with its accessory proteins, TTC7 (two possible isoforms TTC7A and TTC7B) and FAM126 (two possible isoforms FAM126A and FAM126B) forming an ∼750 kDa dimer of heterotrimers, referred to here as the PI4KA complex(7, 8). Both TTC7 and FAM126 contribute towards complex stability, with FAM126 not directly interacting with PI4KA, but instead acting as a stabilizer of TTC7 (9). The PI4KA complex has to be recruited to the PM for its activity (10), primarily through interactions with palmitoylated EFR3 proteins (two isoforms EFR3A and EFR3B). This recruitment of the PI4KA complex to the plasma membrane is mediated by the C-terminus of EFR3 interacting with both TTC7 and FAM126 (11).

Dysregulation of PI4KA and its regulatory partners have been heavily implicated in multiple human diseases (12, 13), with diseases driven by both loss-of-function and gain-of-function alterations. These alterations are often isoform and tissue specific. Loss-of-function mutations in TTC7A are found in patients with immunodeficiencies, intestinal atresia and early onset inflammatory bowel disease (IBD) (14–17) and result in a marked reduction in PI4KA activity. However, there are no known mutations in TTC7B linked to disease. Similarly, disease linked mutations are only found in FAM126A and not in FAM126B. Loss-of-function mutations in FAM126A cause hypomyelination and congenital cataract (HCC), a disorder characterized by neurological impairment, cognitive defects and peripheral neuropathy (18–20). Conversely, overactivation of PI4KA signaling is also involved in disease. In hepatitis C viral infection, the viral protein NS5A activates PI4KA, leading to the formation of viral replication organelles enriched in PI4P (21, 22). Increased PI4KA activity also plays a role in cancer, with overexpression of PI4KA and EFR3A implicated in KRAS driven pancreatic cancers (23, 24). Interestingly, EFR3A interacts with oncogenic KRAS promoting tumorigenic activity (23), with this proposed to be mediated by increased PS levels at the PM (24). The role of PI4KA and EFR3A in driving KRAS driven cancers has renewed enthusiasm in generating pharmacological agents that limit PI4KA signaling in combination with KRAS mutant selective inhibitors.

PI4KA inhibitors were originally developed as anti-hepatitis C viral agents, leading to development of multiple well validated potent and selective PI4KA inhibitors (25, 26). However, enthusiasm for these molecules as therapeutics was greatly diminished as knockdown of PI4KA either genetically or pharmacologically caused rapid death in mice driven by severe gastrointestinal side effects and dramatically decreased PM PIP_2_ levels (26). While there may be possibilities to target PI4KA signaling in EFR3A driven tumors, the severe side effects of ATP-competitive PI4KA inhibitors suggests there could be advantages of novel allosteric PI4KA modulators. The development of molecules that block EFR3 recruitment of PI4KA offers a potential avenue to design novel therapeutics and provides opportunities to further understand how PI4KA is regulated. One of the most well validated tools to manipulate signaling proteins are nanobodies, which bind to specific effector interfaces, leading to modulation of intracellular signaling pathways. Nanobodies are composed of the variable domains (VHH) of heavy chain only antibodies (HCAbs) that lack the light chain of conventional antibodies (27). The single domain architecture results in a compact size (∼15 kDa) that can be readily expressed in bacteria with high yield, making them an attractive tool for therapeutic and diagnostic applications (28).

Understanding regulation of the PI4KA complex requires insight into how different isoforms of its accessory proteins influence plasma membrane recruitment, along with the development of tools to selectively target these isoforms. Here, we have systematically characterized all possible combinations of PI4KA accessory protein isoforms and determined the EFR3A-TTC7B-FAM126A complex as the highest affinity complex. Using a yeast surface display approach, we purified a TTC7B selective nanobody that blocked EFR3 binding, which we named F3IN (E**F**R**3**-**I**nterfering **N**anobody). Using a synergy of cryo-electron microscopy (cryo-EM) and hydrogen-deuterium exchange mass spectrometry (HDX-MS), we identified the binding epitopes of this nanobody on PI4KA, TTC7B and FAM126A and the conformational dynamics upon nanobody binding. We also provide evidence that the nanobody decreases membrane recruitment of PI4KA both on lipid bilayers and in cells, and decreases PM PI4P levels. Overall, this study provides proof-of-concept that PI4KA activity can be modulated with molecules outside the ATP-binding site and introduces a useful isoform-specific nanobody tool to dissect the functional roles of PI4KA accessory proteins in controlling PI4KA activity.

## Results

### EFR3A binds with highest affinity towards TTC7B-FAM126A

The structure of EFR3A bound to the complex of PI4KA with TTC7B and FAM126A suggested that there may be differential affinities of unique EFR3 isoforms to different TTC7 and FAM126 isoforms. This was based on the C-terminal interacting region of EFR3 binding to both TTC7 and FAM126 (11), with the interacting regions in EFR3, TTC7 and FAM126 being highly conserved, but not strictly conserved between isoforms. The different isoforms of EFR3, TTC7, and FAM126 exhibit tissue-specific expression patterns, suggesting distinct physiological roles. Although there are eight possible unique EFR3-TTC7-FAM126 complexes, it remains unclear whether they differ in association. To investigate potential affinity differences between isoform complexes, we purified different combinations of TTC7 and FAM126, as well as MBP tagged C-terminal fragments of EFR3A (721-791) and EFR3B (716-787) (**Fig 1A**). The dimers of TTC7B with FAM126A (1-308) or FAM126B (1-308) were purified from *E. coli,* with the dimers of TTC7A with FAM126A/B purified from *Sf9* cells. All four dimers eluted from gel filtration at sizes consistent with a heterodimer.

**Figure 1.**
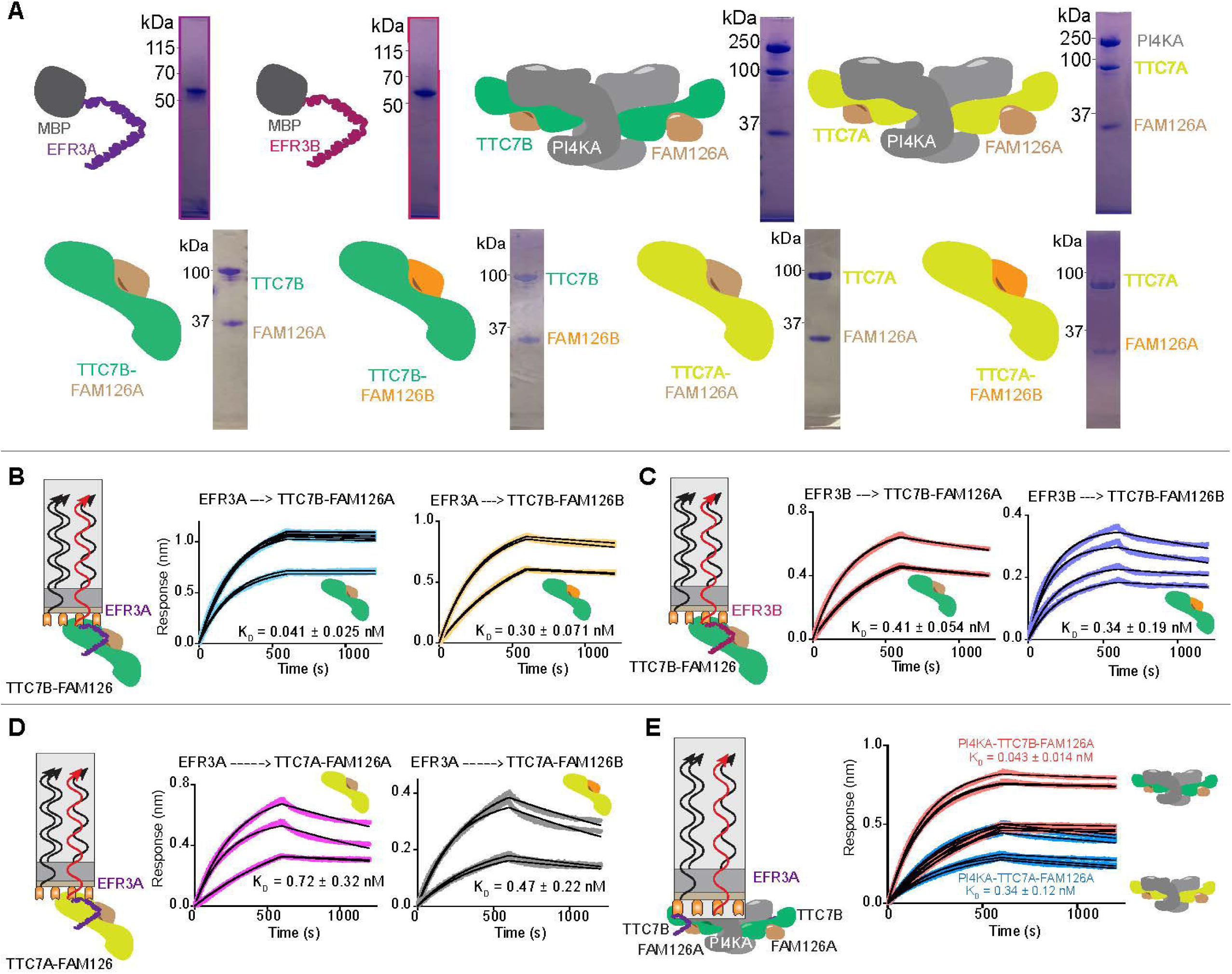
Isoform differences in EFR3. (A) Cartoon representations of the different isoforms of EFR3, TTC7 and FAM126 used in this paper. (L-R) MBP-EFR3A (721-791), MBP-EFR3B (716-787), PI4KA-TTC7B-FAM126A (1-308), PI4KA-TTC7A-FAM126A (1-308), TTC7B-FAM126A (1-308), TTC7B-FAM126B (1-308), TTC7A-FAM126A (1-308), and TTC7B-FAM126B (1-308). SDS gel showing purity are shown, with uncropped gels available in the source data. (B) (L) Cartoon of BLI experiment with EFR3A on the tip and TTC7B-FAM126A/B in solution. Raw BLI traces of EFR3A binding to TTC7B-FAM126A/B. The K_D_ was calculated using a 1:1 binding model (n=4). (C) (L) Cartoon of BLI experiment with EFR3B on the tip and TTC7B-FAM126A/B in solution. Raw BLI traces of EFR3B binding to TTC7B-FAM126A/B. The K_D_ was calculated using a 1:1 binding model (n=4). (D) (L) Cartoon of BLI experiment with EFR3A on the tip and TTC7A-FAM126A/B in solution. Raw BLI traces of EFR3A binding to TTC7A-FAM126A/B. The K_D_ was calculated using a 1:1 binding model (n=4). (E) (L) Cartoon of BLI experiment with EFR3A on the tip and PI4KA complex in solution. (R) Raw BLI traces of EFR3A binding with PI4KA-TTC7A/B-FAM126A (2.5 nM, 5 nM). The K_D_ was calculated using a 1:1 binding model (n=3).

We performed bio-layer interferometry (BLI) experiments to measure binding kinetics with His-tagged MBP-EFR3 immobilized on the biosensor tip and TTC7-FAM126 as the analyte. For all measurements we used MBP alone as a non-specific binding control. The full binding parameters (K_d_, k_on_ and k_off_) are shown in Table S1. Most of the isoform combinations generated BLI traces that fit well to a 1:1 binding model (**Fig 1B-D**), with the only exception being EFR3B binding to TTC7A with either FAM126 isoform (Fig S1). All heterodimers of TTC7 and FAM126 had dissociation constants indicative of relatively similar affinities (∼0.3 nM), with a single exception being EFR3A, which showed almost a 10-fold increase in affinity to TTC7B-FAM126A (0.041 ± 0.025 nM) **(Fig. 1B)**. This increased affinity of EFR3A for TTC7B-FAM126A was primarily driven by a decrease in the k_off_ rate for EFR3A-TTC7B-FAM126A.

EFR3 isoforms primarily recruit TTC7 and FAM126 in the biologically relevant complex also containing PI4KA, with each complex containing two possible EFR3 binding sites due to PI4KA dimerization. We purified heterotrimers of PI4KA with either TTC7A-FAM126A or TTC7B-FAM126A and tested their binding to the C-terminal fragment of EFR3A. The binding affinity of EFR3A to the PI4KA-TTC7B-FAM126A (0.043 ± 0.014 nM ) and PI4KA-TTC7A-FAM126A (0.34 ± 0.13 nM) trimers was similar to that of the dimers, further supporting that TTC7B binds to EFR3A more strongly than TTC7A. **(Fig. 1E)**. Overall, these results show that *in vitro,* the highest affinity interaction between possible EFR3, TTC7, and FAM126 isoforms occurs with the combination of EFR3A and TTC7B-FAM126A.

### EFR3 recruitment to TTC7-FAM126 can be altered by phosphorylation

Alterations in the affinity of EFR3 isoforms binding TTC7-FAM126 led us to interrogate possible differences between them. We first analyzed the EFR3 C-terminus for potential changes in post-translational modifications that may regulate PI4KA recruitment. In yeast, phosphorylation of Efr3 weakens its interaction with Ypp1 (homolog of TTC7) and inhibits the recruitment of Stt4 (homolog of PI4KA) to the membrane (29). There are several reported EFR3A and EFR3B phosphorylation sites that are located at the interface with TTC7 and FAM126, including Y723 in EFR3B and S738 in EFR3A (30). Y723 in EFR3B lies at the predicted interface with TTC7, where its phosphorylation may disrupt binding due to the proximity of the hydroxyl group to a conserved glutamic acid residue in both TTC7A and TTC7B **(Fig. 2A)**. We used kinase motif analysis to identify possible kinases that may phosphorylate Y723, with it being identified as a likely substrate of multiple Src family kinases (top five scoring kinases: BLK, HCK, FRK, LYN, and LCK) (31). We treated EFR3B with an active construct of the tyrosine kinase LCK, which resulted in predominantly singly phosphorylated species at pY723 (∼92%) as determined by MS analysis **(Fig. 2B-C)**. We next performed BLI to measure the binding of phospho-EFR3B to TTC7B-FAM126A and observed a ∼90% decrease in binding upon Y723 phosphorylation **(Fig. 2D)**. These results demonstrate that phosphorylation of specific EFR3 isoforms may provide another modality to regulate isoform-specific interactions and PI4KA recruitment and activation at the PM.

**Figure 2.**
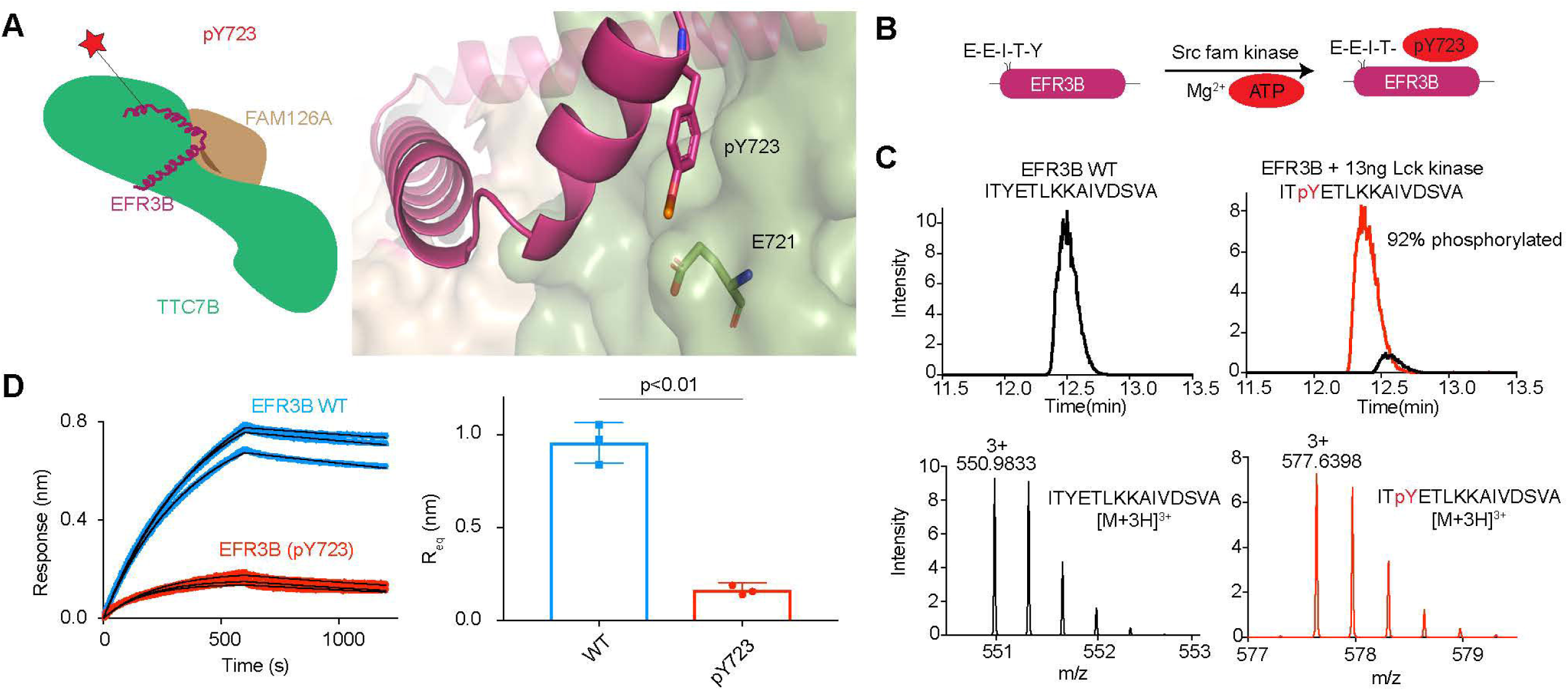
EFR3B phosphorylation decreases TTC7-FAM126 binding. (A) (L) Cartoon schematic of EFR3B-TTC7B-FAM126A with Y723 highlighted with a red star. (R) AlphaFold3 model of EFR3B-TTC7B-FAM126A showing phosphorylated Y723 on EFR3B and E721 on TTC7B. (B) Schematic of EFR3B phosphorylation by Src family kinases. (C) (Top) Extracted Ion Chromatograms (EIC) of the EFR3B peptide for unphosphorylated (Black) and phosphorylated (Red). (Bottom) Peptide spectra of the unphosphorylated (L) and phosphorylated (R) EFR3B peptide with Y723 highlighted red in the sequence. (D) Raw BLI traces of EFR3B WT (blue) or pY723 (red) binding to TTC7B-FAM126A (L). Maximum BLI response of EFR3B WT and pY723 (R). Error is shown as SD (n = 3) with p value indicated.

### Screening of putative inhibitors of PI4KA

Extensive biochemical screening of small molecule libraries resulted in the development of multiple potent, and highly selective PI4KA inhibitors, although early testing in mouse models revealed severe toxicity (25, 26). More recently, however, it has been proposed that there may be a therapeutic window when these compounds are used in combination with RAS-targeting inhibitors (23, 24). This prompted us to determine whether the two TTC7 isoforms differentially affect complex inhibition in PI4KA-TTC7A/B-FAM126A assemblies. We also tested the FDA approved molecule simeprevir, which has been putatively described as a PI4KA inhibitor (24, 32), although, importantly the published *in vitro* inhibition data did not seem to show profiles consistent with ATP competitive inhibition (32).

PI4KA kinase assays were carried out with the three chemical modulators, GSK-A1, GSK-F1, and simeprevir for both PI4KA-TTC7A-FAM126A and PI4KA-TTC7B-FAM126A in the presence of lipid vesicles composed of 100% PI. Both complexes were inhibited by the GSK-A1 and GSK-F1 compounds, however, we detected no inhibition by simeprevir at concentrations up to 10 µM (**Fig 3**). These results indicate that the two TTC7 isoforms do not differentially affect PI4KA complex inhibition by ATP-competitive inhibitors and suggest that previous reports proposing simeprevir as a PI4KA inhibitor should be interpreted with caution.

**Figure 3.**
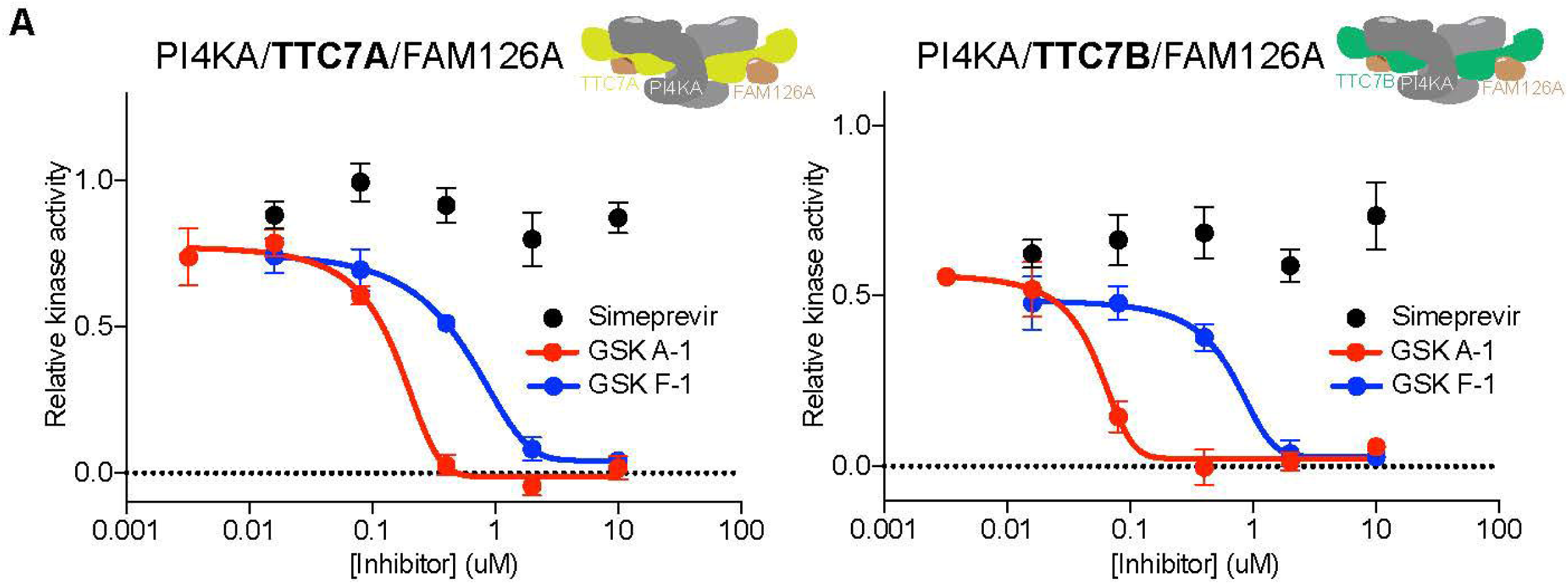
Simeprevir does not inhibit PI4KA activity. (A) IC_50_ curves of (L) PI4KA/TTC7A/FAM126A and (R) PI4KA/TTC7B/FAM126A when treated with increasing concentrations of GSK-A1 (red) and GSK-F1 (blue), compared to PI4KA activity when treated with increasing concentrations of simeprevir (black). Assays were carried out with 100% PI vesicles at a final concentration of 0.25 mg/ml in the presence of 100 uM ATP. PI4KA concentration was 5 nM. Error is shown as standard deviation (n=3).

### Generation of a nanobody that blocks EFR3 recruitment of TTC7B-FAM126A

As the TTC7B-FAM126A isoform showed the highest affinity for EFR3 recruitment, and PI4KA inhibitors showed no differences in inhibition between PI4KA complexes, we sought to develop a tool for selectively targeting unique PI4KA complexes. To address this, we aimed to isolate a nanobody that specifically blocked EFR3 binding to the TTC7B-FAM126A complex. For this purpose, we screened a yeast surface display synthetic library of >5 x 10^8^ nanobodies for binders to the EFR3 binding site of TTC7B-FAM126A (33). We used the TTC7B-FAM126A (1–308) heterodimer as our antigen **(Fig. 4A)** and performed multiple rounds of magnetic-activated cell sorting (MACS) and fluorescence-activated cell sorting (FACS) to obtain a library of nanobodies that bound TTC7B-FAM126A **(Fig. 4B)**. To specifically obtain nanobodies that bind at the EFR3 binding site on TTC7B-FAM126A, we purified TTC7B in complex with a chimeric fusion construct comprising FAM126A (1-308) fused to the C-terminus of EFR3A (681-791). The gel filtration elution profile indicated the formation of a heterodimer **(Fig. 4A+C)**. We used this construct to counter-select against our original nanobody library **(Fig. 4B)**, resulting in the isolation of a nanobody that bound TTC7B-FAM126A but not the TTC7B-EFR3A-FAM126A chimera. We named this nanobody, F3IN (E**F**R**3**-**I**nterfering **N**anobody). BLI confirmed that F3IN binds TTC7B-FAM126A with high affinity, while showing significantly reduced binding to the TTC7B-EFR3A-FAM126A chimera **(Fig. 4D)**.

**Figure 4.**
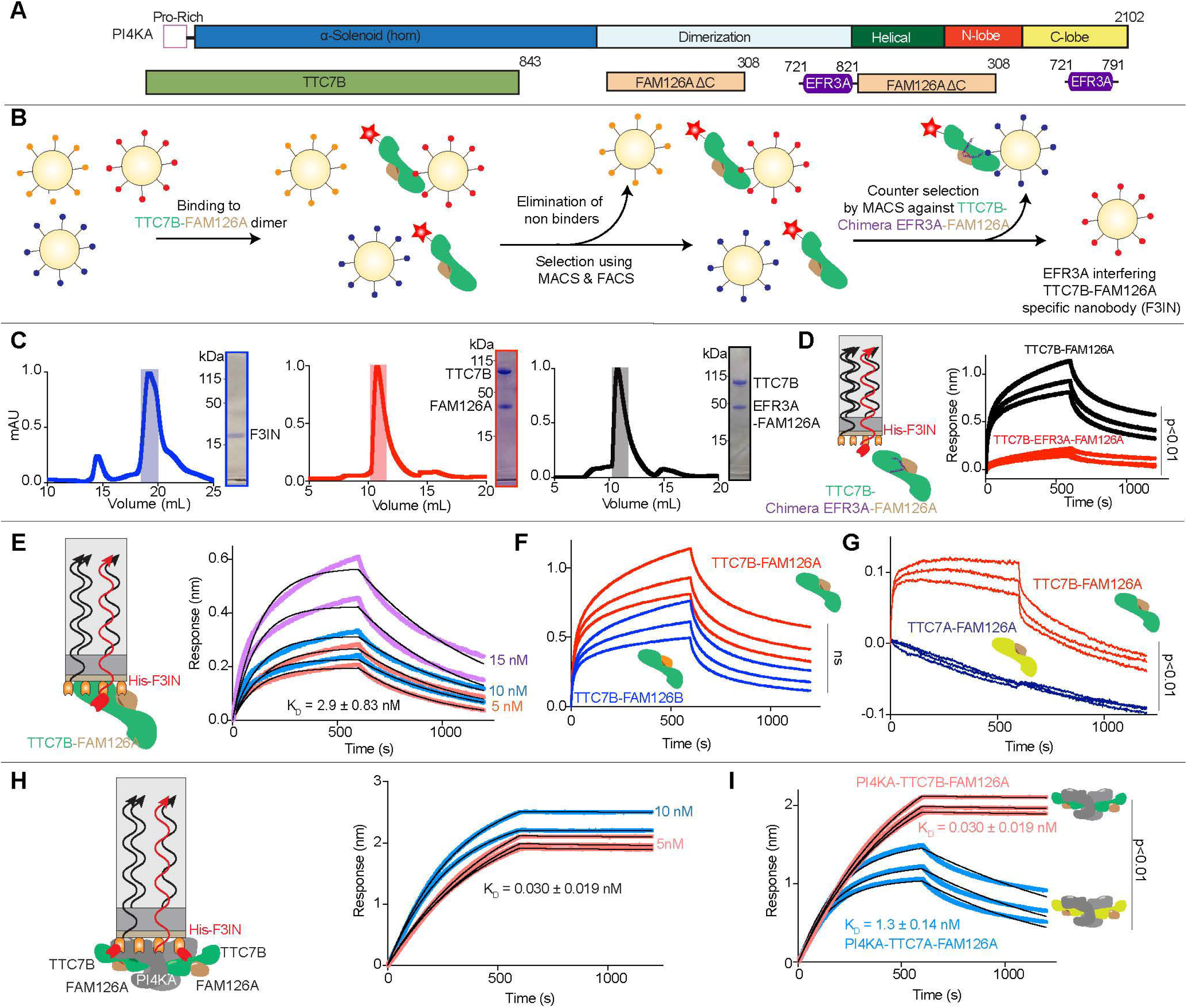
TTC7B selective nanobody generation using yeast surface display. (A) Domain schematics of PI4KA-TTC7B-FAM126A (1-308) (referred to as PI4KA complex), EFR3A (721-791) and Chimeric EFR3A-FAM126A (1-308) constructs used in this paper. (B) Schematic of the nanobody selection process using yeast surface display. Antigen is shown with a fluorescent tag (red star). Nanobodies with affinity to antigen are selected using MACS and FACS. Counter selection was performed against chimeric TTC7B-EF3A-FAM126A using MACS to obtain TTC7B-FAM126A specific nanobody (named F3IN). (C) Size exclusion chromatography traces and corresponding SDS-PAGE gels of F3IN, TTC7B-FAM126A and chimeric TTC7B-EFR3A-FAM126A. (D) Cartoon of BLI experiment with F3IN on the tip and TTC7B-FAM126A or TTC7B-chimera EFR3A-FAM126A in solution. BLI traces of F3IN against TTC7B-FAM126A and chimeric TTC7B-EFR3A-FAM126A. (E) Cartoon of BLI experiment with F3IN on the tip and TTC7B-FAM126A in solution. Raw BLI traces of F3IN (100 nM) binding to TTC7B-FAM126A (5 nM, 10 nM, 15 nM). The K_D_ was calculated using a 1:1 binding model (n=4). (F) Raw BLI traces for the binding of F3IN (100 nM) to TTC7B-FAM126A (50 nM) and TTC7B-FAM126B (15 nM). (G) Raw BLI traces for the binding of F3IN (100 nM) to TTC7B-FAM126A (15 nM) and TTC7A-FAM126A (15 nM). (H) Cartoon (L) of the BLI experiment. Raw BLI traces of F3IN (100 nM) binding to PI4KA-TTC7B-FAM126A (2.5 nM, 5 nM, 10 nM). The K_D_ was calculated using a 1:1 binding model (n=3). (I) Raw BLI traces for the binding of F3IN (100 nM) to PI4KA-TTC7B-FAM126A (5 nM) and PI4KA-TTC7A-FAM126A (5 nM). The K_D_ was calculated using a 1:1 binding model (n=3).

The F3IN nanobody bound to the TTC7B-FAM126A dimer with a K_D_ of 2.9 ± 0.83 nM **(Fig. 4E)** and showed similar binding to dimers of TTC7B-FAM126B **(Fig. 4F**, Full kinetic parameters of F3IN binding are in table S1). In contrast, F3IN showed no detectable binding to TTC7A-FAM126A dimers, indicating isoform selectivity towards TTC7B **(Fig. 4G).** The most physiologically relevant binding interaction of F3IN would be with the PI4KA complex, therefore we performed BLI experiments with PI4KA trimer in solution. Surprisingly, we found that the F3IN nanobody bound with a ∼100-fold increased affinity to the PI4KA-TTC7B-FAM126A complex compared to the TTC7B-FAM126A dimer (K_D_ of 0.030 ± 0.019 nM to the trimer) **(Fig. 4H)**. In alignment with dimer results, we observed a ∼30-fold decrease in affinity of F3IN for PI4KA-TTC7A-FAM126A (0.030 nM for TTC7B versus 1.3 nM for TTC7A complex). Overall, this showed that the F3IN nanobody is specific for TTC7B over TTC7A and binds with higher affinity to the PI4KA heterotrimer over the TTC7-FAM126 dimer.

### Cryo-EM analysis of F3IN bound to PI4KA-TTC7B-FAM126A

To identify the binding interface of F3IN with the PI4KA complex, we generated a 3D reconstruction of the PI4KA-TTC7B-FAM126A-F3IN nanobody complex, using cryo-EM, at a nominal resolution of 3.54 Å (from 282,982 particles **(Fig 5A-B, S2A-F, Table S2)**. Samples for cryo-EM analysis were prepared by brief treatment with a crosslinker to stabilize the PI4KA complex, similar to our previous cryo-EM analysis of PI4KA regulatory complexes (11, 34). Local masked refinement centered on the F3IN-TTC7B interface enabled unambiguous determination of the residues mediating the interaction between F3IN and the PI4KA complex. Using a combination of manual model building and AlphaFold3 (35), we were able to generate a molecular model of F3IN binding to the PI4KA-TTC7B-FAM126A trimer with the density allowing for manual building of the CDR loops mediating interaction with the trimer **(Fig S2G)**. The structure revealed that F3IN included an additional extended beta strand within the CDR3 loop, which orients it into the cavity normally occupied by the α3 helix of EFR3A **(Fig 5C-D)**. This orientation suggests that F3IN sterically blocks EFR3A binding by occluding its binding with the CDR3 loop.

**Figure 5.**
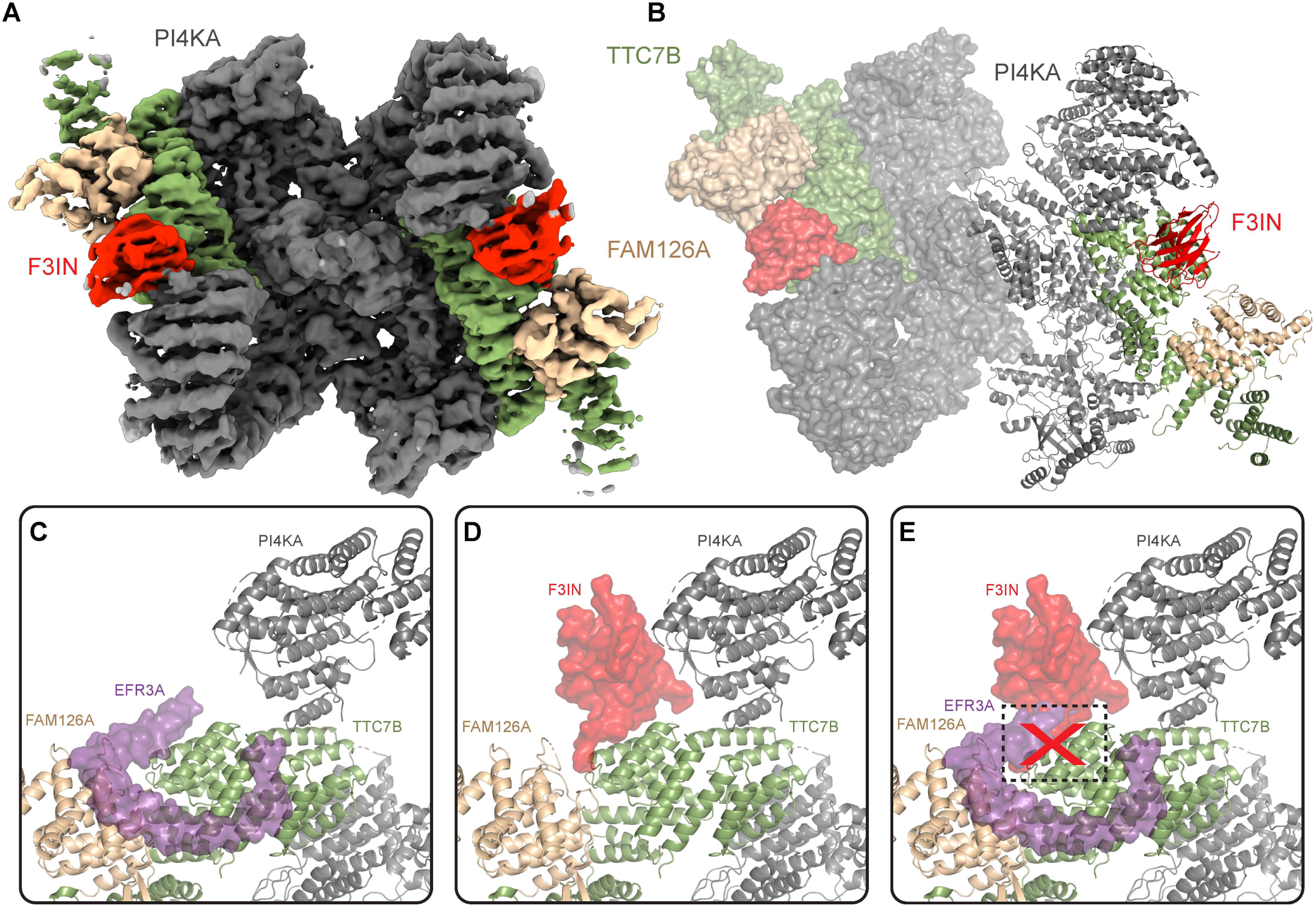
Cryo-EM analysis of F3IN binding to the PI4KA complex. (A) Cryo-EM density map of the PI4KA complex bound to F3IN. (B) Molecular model of F3IN bound at the interface of TTC7B and the PI4KA horn domain. (C) Zoomed in cartoon of the C-terminus of EFR3A bound to the PI4KA complex. (PDB: 9BAX) (D) Zoomed in cartoon of F3IN bound at the interface of TTC7B and PI4KA. (E) Overlay of the PI4KA-EFR3A (PDB: 9BAX) and PI4KA-F3IN models depicting binding of F3IN and EFR3A at the same TTC7B helix (represented as a black box with a red cross).

The TTC7B-F3IN interface is composed of 415 Å^2^ buried surface area. The interface is mainly mediated by three TTC7B α helices, spanning residues 535-545, 546-563, and 564-578. These helices interact directly with the extended β-strand in CDR3 of F3IN. Although TTC7A and TTC7B have a sequence similarity of 49.7% with a highly conserved C-terminus, our structure revealed possible insight for why F3IN was selective to TTC7B containing heterodimers. The most prominent difference is a glycine residue in TTC7B (G546) that is not conserved in TTC7A, where it is replaced by a lysine (K563). In the cryo-EM structure, G546 lies in close spatial proximity to R105 in the CDR3 of F3IN. This positioning suggests that the corresponding lysine in the TTC7A isoform would result in a charge-charge repulsion with R105, potentially preventing nanobody binding. Intriguingly, F3IN also interacts with the N-terminus of PI4KA resulting in 280 Å^2^ of buried surface area. This interface is mediated by the N-terminus of F3IN and the N-terminal horn of PI4KA, comprising a β-strand (residues 67-73) and a loop (residues 58-66). This additional interface likely explains the increased affinity of F3IN with the PI4KA complex, and why there was weak affinity for the PI4KA-TTC7A-FAM126A complex, but not the TTC7A-FAM126A dimer.

### HDX-MS validation of epitopes on PI4KA and TTC7B

To further understand the dynamics of the interaction between PI4KA and F3IN, we used HDX-MS on the PI4KA complex and PI4KA-F3IN complex. HDX-MS is a powerful technique to identify binding epitopes and investigate dynamic conformational changes. It measures the exchange of backbone amide hydrogens with deuterium (36, 37), with the rate of exchange being primarily dependent on secondary structure (38). HDX-MS experiments were performed comparing the PI4KA-TTC7B-FAM126A complex alone and PI4KA-TTC7B-FAM126A complex bound to F3IN. We incubated the samples with D_2_O buffer at 4 different timepoints (3s, 30s, 300s, 3000s at 18°C) with significant differences between the two conditions defined as changes >0.4 Da, 5% at any given time point and an unpaired t-test value of p<0.01. Upon F3IN binding, we observed a significant decrease in deuterium incorporation at the N-terminus of the PI4KA horn spanning residues 51-83 **(Fig. 6A-C)**, showing a clear epitope on PI4KA and supporting our cryo-EM model. Surprisingly, we saw no clear decrease in exchange on TTC7B and instead observed an increase in HDX at residues 539-545 in TTC7B.

**Figure 6.**
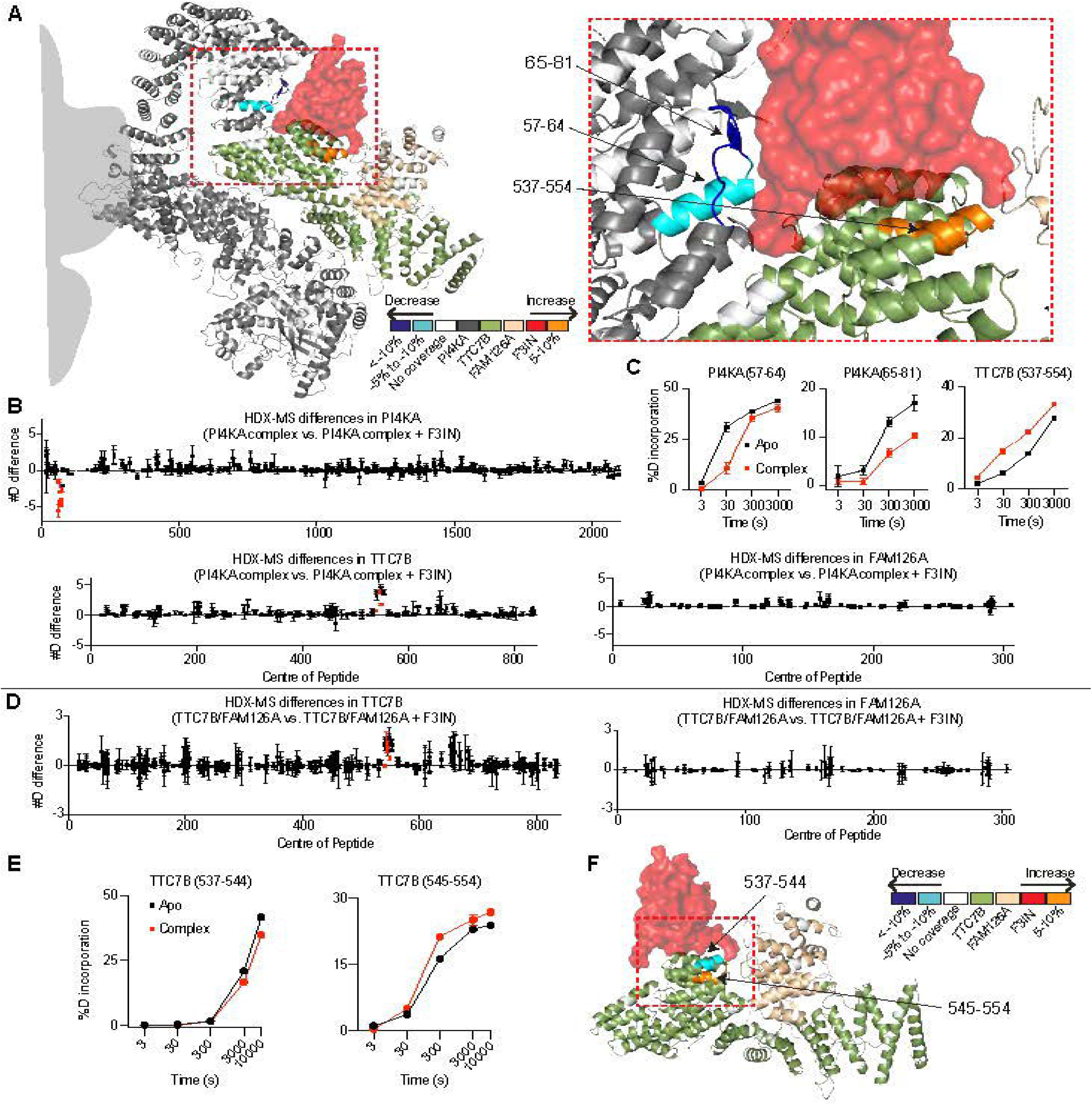
HDX-MS analysis of the interaction of F3IN with PI4KA and TTC7B. (A) HDX differences (significance defined as >5% and 0.4 Da with P< 0.01 in an unpaired two-tailed t test at any time point) upon PI4KA complex binding to F3IN mapped on PI4KA-F3IN structure. (B) Sum of the deuteron difference of the PI4KA complex upon binding to F3IN across the entire time course. Each point represents a single peptide. Peptides that met the significance criteria described in (A) are colored red. Error is shown as the sum of SDs across all time points (n=3). (C) Selected deuterium exchange time courses of PI4KA peptides showing significant changes (significance criteria in A) at any timepoint in the presence (red) and absence (black) of F3IN. (D) Sum of the deuteron difference of TTC7B-FAM126A upon binding to F3IN across the entire time course. Peptides that met the significance criteria described in (A) are colored red. Error is shown as the sum of SDs across all time points (n=3). (E) Selected deuterium exchange time courses of TTC7B peptides showing significant changes (significance criteria in A) at any timepoint in the presence (red) and absence (black) of F3IN. (F) HDX differences upon TTC7B-FAM126A binding to F3IN mapped on PI4KA-F3IN structure.

In the absence of PI4KA, the TTC7B-FAM126A dimer also showed F3IN binding with high affinity, therefore we used HDX-MS to map the epitope and investigate the differences in conformational dynamics of TTC7B-FAM126A upon F3IN binding. HDX-MS experiments were performed to compare TTC7B-FAM126A in the presence and absence of F3IN at 5 different timepoints (3s, 30s, 300s, 3000s, 10,000s at 18°C). We observed a significant decrease in TTC7B in the peptide 539-544 consistent with the interface observed in our cryo-EM analysis. We also measured a significant increase in HDX in the subsequent peptide 545-554 **(Fig. 6D-F)**. Overall, this suggests that nanobody binding induces a shift in dynamics in TTC7B, which may provide another mechanism for selectivity compared to TTC7A.

### Nanobody F3IN competitively inhibits EFR3A binding

The cryo-EM structure and HDX-MS results indicated that F3IN forms a dual interface with both TTC7B and PI4KA. To investigate whether F3IN is a competitive inhibitor of EFR3 for the TTC7B binding site, we developed a competition BLI assay. Here, F3IN was loaded onto the biosensor and dipped into solution containing either 50 nM of TTC7B-FAM126A or 50 nM of TTC7B-FAM126A with increasing concentrations of EFR3A (15 nM to 240 nM). We found that EFR3A inhibited the binding of F3IN to TTC7B-FAM126A **(Fig. 7A)**. This indicates that both EFR3A and F3IN compete for the same binding site on TTC7B in agreement with the cryo-EM and HDX-MS results.

**Figure 7.**
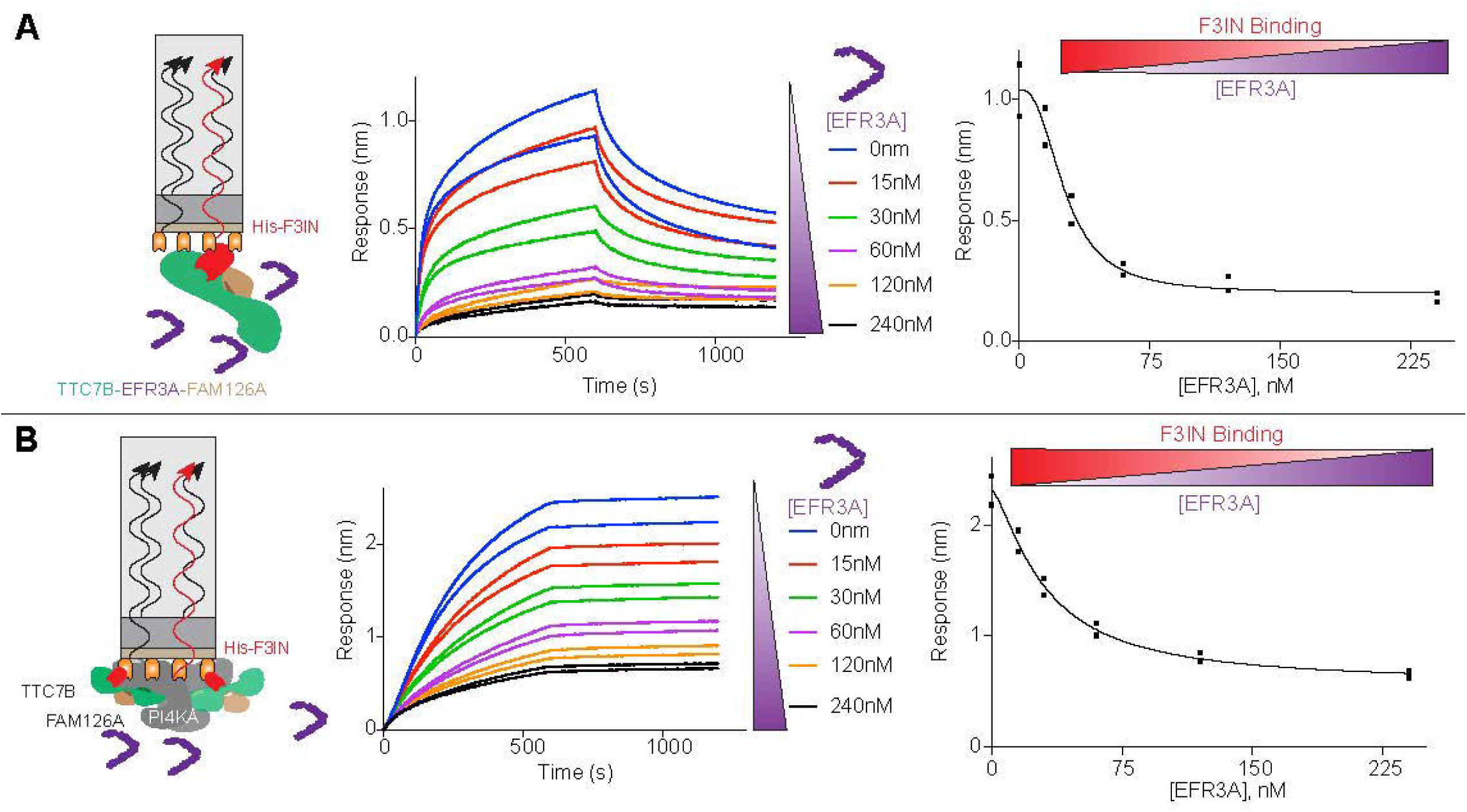
EFR3A competitively inhibits binding to F3IN for both the TTC7B-FAM126A dimer and the PI4KA-TTC7B-FAM126A trimer. (A) Cartoon (L) of BLI experiment with immobilized F3IN dipping into TTC7B-FAM126A solution with increasing concentrations of EFR3A. Raw association and dissociation curves (M) for the binding of F3IN (100 nM) to TTC7B-FAM126A (50 nM) at varying concentrations of EFR3A. (R) Dose response curve showing decrease in F3IN binding upon increase in EFR3A concentrations. (B) Cartoon (L) of BLI experiment with immobilized F3IN dipping into PI4KA complex solution with increasing concentrations of EFR3A. Raw association and dissociation curves (M) for the binding of F3IN (100 nM) to PI4KA complex (10 nM) at varying concentrations of EFR3A. (R) Dose response curve showing decrease in F3IN binding upon increase in EFR3A concentration.

However, due to the unexpected interaction of F3IN with the horn region of PI4KA, it was important to perform a BLI competition assay with the full PI4KA complex to determine the degree of inhibition with the physiologically relevant complex. In this assay, we used the same above-mentioned conditions with 10 nM PI4KA rather than the TTC7B-FAM126A dimer. As expected, we observed a decrease in F3IN-PI4KA binding with increasing concentration of EFR3A **(Fig. 7B)**. Altogether, these results indicate that the F3IN nanobody competes with EFR3A for binding to TTC7B and further validates its usefulness as a tool to specifically inhibit TTC7B containing complexes of PI4KA.

### Nanobody F3IN blocks EFR3 mediated membrane recruitment and decreases EFR3 mediated plasma membrane localization

To examine if the F3IN nanobody can block EFR3A mediated membrane recruitment of PI4KA we performed single molecule TIRF microscopy experiments on supported lipid bilayers (SLBs). We tethered EFR3A to SLBs using the SpyCatcher-SpyTag system (39) (**Fig. 8A**). Membranes containing cysteine-reactive maleimide lipids were used to covalently attach a cys labelled SpyCatcher to lipid bilayers. We then recruited EFR3A to these membranes by using a construct containing a N-terminal GFP-SpyTag-fused to residues 721-791 of EFR3A (referred to as GFP-SpyTag-EFR3A). We visualized the membrane recruitment of AF647 labelled PI4KA-TTC7B-FAM126A (AF647-PI4KA) in the absence and presence of EFR3A (**Fig. 8B**). AF647-PI4KA was only robustly recruited to the membrane when EFR3A was present on the SLBs. Addition of the F3IN nanobody to EFR3A containing membranes dramatically reduced the single molecule membrane binding frequency (**Fig. 8C**) and bulk membrane recruitment (**Fig. 8D**) of AF647-PI4KA in a dose dependent manner (**Fig. 8E**).

**Figure 8.**
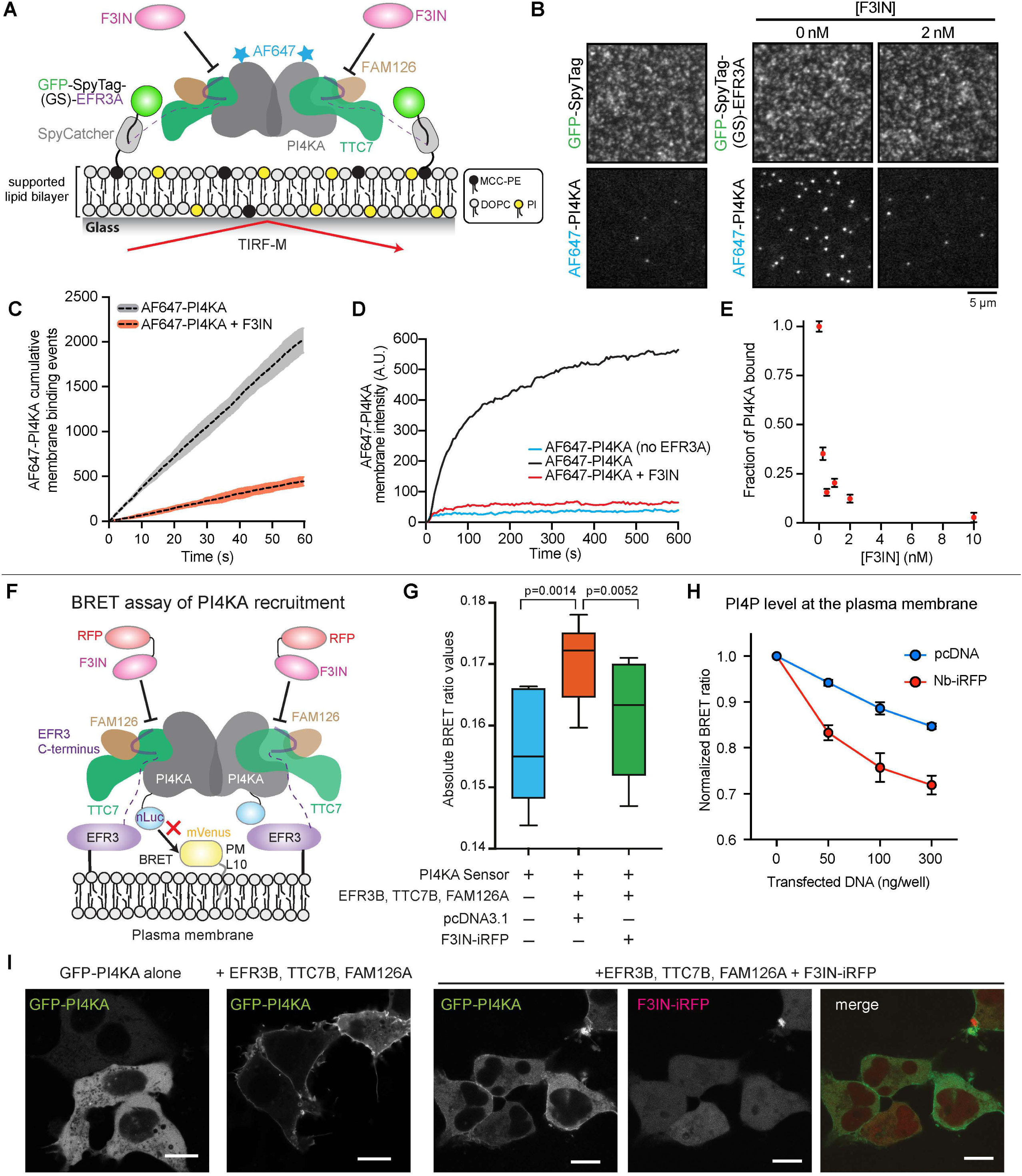
F3IN blocks EFR3 mediated PI4KA membrane recruitment on both supported lipid bilayers and in HEK293 cells. (A) Cartoon schematic showing the experimental design for measuring AF647-PI4KA localization on a SLBs containing tethered EGFP-SpyTag-EFR3A. SpyCatcher is covalently linked to a maleimide lipid (i.e. MCC-PE), while EGFP-SpyTag-EFR3A is conjugated to SpyCatcher via an isopeptide bond. Localization of EGFP-SpyTag-EFR3A and AF647-PI4KA is visualized by TIRF microscopy. (B) Representative TIRF microscopy images showing the localization of membrane-tethered EGFP-SpyTag-EFR3A and EGFP-SpyTag. Localization of 1 nM AF647-PI4KA was visualized in the absence and presence of 2 nM nanobody, F3IN. (C) The single molecule membrane binding frequency of 1 nM AF647-PI4KA is reduced in the presence of 2 nM F3IN. Total number of membrane binding events were measured on a 3000 µm^2^ membrane surface area by smTIRF microscopy. Curves represent the mean (dashed black line) and standard deviation (n=3). (D) Kinetic traces showing the bulk membrane absorption and equilibration of 1 nM AF647-PI4KA on EGFP-SpyTag-EFR3A (or EGFP-SpyTag) membranes in the absence and presence of 1 nM nanobody, F3IN. (E) Nanobody, F3IN, reduces membrane localization of 1 nM AF647-PI4KA in dose dependent manner. Plot shows the normalized AF647-PI4KA membrane fluorescence as a function of F3IN concentration. (B-E) Membrane composition: 94% DOPC, 4% PI, 2% MCC-PE (SpyCatcher conjugated). (F) Cartoon depicting the quantitative BRET-based PI4KA recruitment assay. Briefly, the PM-anchored BRET acceptor (L10-mVenus) is located near the nLuc-tagged PI4KA BRET-donor, and thereby efficiently increase the relative BRET signal, only if co-assembled with the requisite molecular partners for PM targeting. Co-expression of the nanobody-iRFP construct reduces BRET signal by blocking membrane localisation of PI4KA. (G) Normalized BRET signal measured from HEK293A cell populations (∼0.5 x 10^5^ cells/well) expressing a fixed amount of the PM-PI4KA^BRET^ biosensor (L10-mVenus-tPT2A-NLuc-PI4KA; 500 ng/ well) together with 300 ng/well of the indicated constructs. (BRET sensor alone (blue) or together with EFR3B-P2A-TTC7B-T2A-FAM126A and pcDNA3.1 (red) or with EFR3B-P2A-TTC7B-T2A-FAM126A and F3IN-iRFP (green). Absolute BRET values are shown from 6 independent experiments each performed in triplicates. One Way ANOVA with Geisser-Greenhouse correction was used to compare the various groups with multiple comparisons, where values obtained from each experiment were treated as matched data. (H) Normalized BRET signal measured from HEK293A cell populations (∼0.5 x 10^5^ cells/well) expressing a fixed amount of the PM-PI(4)P biosensor (L10-mVenus-tPT2A-NLuc-P4M2x; 100 ng/ well) together with the indicated amounts of control pcDNA3.1 (blue) or F3IN-iRFP (red). BRET values were normalized to the control BRET value without the expression of the additional plasmids. Means ± range of two independent experiments are shown each performed in triplicates. (I) Representative confocal images of live HEK293-AT1 cells co-expressing (50 ng/well) of EGFP-PI4KA alone or together with EFR3B-P2A-TTC7B-T2A-FAM126A (100 ng/well), with or without F3IN-iRFP (100 ng/well).

To test how the nanobody alters PI4KA recruitment in live cells we carried out PI4KA PM recruitment assays using a bioluminescence resonance energy transfer (BRET) between a PM-targeted Venus protein and a nano-Luciferase (nLuc)–tagged PI4KA (**Fig. 8F**), as previously described (11). Live-cell experiments were carried out using human embryonic kidney (HEK) 293-AT1 cells (40) expressing a fixed amount of a PM-PI4KA BRET biosensor and a TTC7B-EFR3A-FAM126A plasmid as previously described (11). Co-transfection of an iRFP tagged F3IN nanobody caused a significant decrease in the BRET signal compared to a pcDNA3.1 vector control (**Fig 8G**). We performed experiments to monitor how expression of the F3IN nanobody affects PM PI(4)P levels using our published BRET analysis (41). These data showed that PM PI4P levels were substantially decreased by expression of increasing amounts of F3IN-iRFP conjugates (**Fig. 8H**). In parallel with this BRET-based approach, we also used confocal microscopy for live-cell imaging studies to examine the subcellular localization of an enhanced green fluorescent protein (EGFP)–tagged PI4KA construct after co-expression of EFR3B, TTC7B, and FAM126A with and without the F3IN nanobody (**Fig 8I**). Consistent with our quantitative BRET-based measurements, we saw a decreased localization of EGFP-PI4KA to the PM when the F3IN nanobody was co-expressed.

## Discussion

Dysregulation of PI4KA and its regulatory partners is involved in multiple human diseases, arising from both gain- and loss-of-function alterations in PI4KA and its accessory proteins EFR3A, TTC7A, and FAM126A. The isoforms TTC7A/B, FAM126A/B, and EFR3A/B are widely expressed, however, there are distinct tissues that show enrichment for specific variants. Clinical mutations in several PI4KA accessory proteins have been identified. For example, patient-derived mutations in TTC7A result in severe intestinal developmental disorders and immunological deficiencies (17), and mutations in FAM126A lead to hypomyelination and congenital cataracts (9). However, no diseases are yet known to be caused by mutations in TTC7B and FAM126B. Although the structure of the C-terminus of EFR3A bound to the PI4KA-TTC7B-FAM126A complex (11) revealed the possibility for distinct association between EFR3 with TTC7-FAM126 isoforms, determining the differences in recruitment of different PI4KA complexes by EFR3 to the PM remained important for defining potential isoform-specific roles. Here we find that most EFR3 isoforms bind to most TTC7-FAM126 complexes with similar affinity (∼0.3 nM), with the exception of EFR3A, which binds with a significantly higher affinity as its interaction with TTC7B-FAM126A had a very slow k_off_ rate. This enhanced affinity for one isoform complex may indicate the need for enhanced PI4P synthesis in EFR3A, FAM126A and TTC7B enriched tissues. However, further work will be required to completely understand the implications of this affinity difference on PI4KA signalling. It is important to note slow dissociation rates (k_off_) were observed for all EFR3 complexes with TTC7-FAM126 isoforms, suggesting all are likely to be quite stable, but with enhanced stability for the EFR3A-TTC7B-FAM126A complex,

The C-terminus of EFR3 has 67% sequence identity between EFR3A and EFR3B. There is differential expression of these isoforms, as EFR3A is highly expressed in testis, fibroblasts, and adipocytes, and EFR3B is enriched in the brain and eye, and its expression in fibroblasts and adipocytes is 30- to 100-fold lower than EFR3A (42, 43). This differential expression suggests distinct tissue specific physiological roles for the two isoforms. The N-terminus of the EFR3 proteins can be palmitoylated at cysteine residues, with a palmitoylation code governing the spatial organisation at the PM (44, 45), further diversifying their regulatory potential. Phosphorylation of Efr3 in yeast disrupts Stt4/Ypp1 (orthologs of PI4KA/TTC7) membrane recruitment, and therefore Stt4 activity (29). Similarly, multiple phosphorylation sites map onto mammalian EFR3A and EFR3B at the TTC7-FAM126 interface (43). The location of these sites is divergent between the two isoforms, suggesting other possible isoform-specific regulation of PI4KA complexes. We determined *in vitro* phosphorylation of EFRB-Y723 by the Src family tyrosine kinase, LCK, markedly decreased binding with TTC7B-FAM126A. However, determining whether LCK is the physiological kinase responsible requires further work, whilst also highlighting the need to screen other Src family kinases as potential regulators of EFR3B, and by extension, regulators of PI4KA activity. Interestingly, Y723 in EFR3B is not conserved in EFR3A, which instead contains a phenylalanine at the corresponding position (F728). This divergence suggests a potential mechanism for differential regulation of the PI4KA complex, possibly influenced by the distinct expression patterns and tissue specificity of EFR3 isoforms. There is precedence for this type of regulation as phosphorylation of EFR3A at S738 by serine/threonine kinases may modulate binding to TTC7B-FAM126A, however, its impact on affinity remains unclear. The top ranked kinase for this site is polo like kinase 1 (PLK1), which plays an important role in regulating cell division. Therefore, phosphorylation of EFR3A may similarly play an important role in controlling PI4KA membrane recruitment and activity. This together with previously identified roles of FAM126 phosphorylation in regulating PI4KA activity (46), provides unique insight into how PTMs can regulate PI4KA activity. Further investigation on how EFR3 phosphorylation is controlled in cells by the specific kinases and phosphatases will be key to understanding PI4KA regulation.

Although the TTC7 isoforms are broadly expressed in mammals, there are key differences. While TTC7B is expressed predominantly in the brain, TTC7A is highly expressed in the gut and thymus with loss-of-function mutations in TTC7A causing severe gastrointestinal diseases (17). Many of these mutations, including S539L, A551D, and H570R, reside in helices that either bind the α3-helix of EFR3A or pack against the interfacial helices. These diseases, caused by lack of PI4KA function, shed insight on why PI4KA inhibitors have severe toxicity in animal models (26). Recently, it has been proposed that PI4KA inhibitors may have a therapeutic window for treating some cancers when used in combination with PI3K- or RAS-targeting inhibitors (23, 24). We demonstrated that both PI4KA complexes are equally sensitive to established PI4KA inhibitors while simeprevir, which was previously reported as a direct inhibitor of PI4KA (24, 32), showed no inhibitory activity even at high concentrations. While opportunities may still exist for using PI4KA inhibitors, the development of isoform-specific biomolecules offer a more refined approach. We reason that blocking EFR3 mediated plasma membrane recruitment of PI4KA could represent a useful approach to regulate its activity. Indeed, modulating PI4KA activity, without fully inhibiting it, could potentially reveal a novel therapeutic strategy.

In recent years, nanobodies have emerged as tools in stabilizing and disrupting protein-protein interactions (47). For example, nanobodies binding to the spike protein of SARS-CoV2 alter its conformational dynamics disrupting binding with host receptors (48). Similarly, in the context of RAS signaling, nanobodies bind to SOS1 and competitively inhibits SOS1-RAS signaling (49). Therefore, there is sufficient evidence for the use of the nanobodies as complex disruptors. Using a yeast display approach, we isolated and characterized a nanobody we called F3IN that binds specifically to TTC7B and competitively inhibits the EFR3-TTC7B interaction. Our cryo-EM and HDX-MS analyses showed that F3IN binds at the interface of PI4KA and TTC7B and induced a conformational change in TTC7B upon binding. We found that the F3IN nanobody could decrease EFR3A mediated PI4KA membrane recruitment both *in vitro* on supported lipid bilayers using TIRF microscopy, and in live cell assays using BRET analysis and confocal microscopy. Moreover, expression of the F3IN nanobody also decreased PI(4)P levels in the PM of intact HEK293-AT1cells. The specificity of F3IN for TTC7B is of particular importance in TTC7B-enriched tissues, as EFR3A has stronger affinity for this isoform compared to TTC7A. From a physiological perspective, the ability to selectively inhibit EFR3 mediated PI4KA recruitment will provide additional insight on PI4KA regulation. Future experiments will need to address what the functional role of blocking EFR3 mediated PI4KA recruitment in specific tissues *in vivo*.

In conclusion, our study provides a detailed molecular framework for understanding isoform-specific regulation of PI4KA and introduces a nanobody-based approach to selectively modulate PI4KA membrane recruitment and activity. These findings lay the foundation for future efforts aimed at the development of more targeted, less toxic strategies to modulate PI4KA function in disease.

## Experimental procedures

### Plasmids

Plasmids containing genes for full-length SSH-PI4KA-TTC7B-FAM126A (1-308) were cloned into the MultiBac vector, pACEBac1 (Geneva Biotech) as described in (8). Plasmids containing genes for full-length SSH-PI4KA-TTC7A-FAM126A (1–308), SSH-ybbr-PI4KA-TTC7B-FAM126A (1-308), SSH-TTC7A-FAM126A (1-308), and SSH-TTC7A-FAM126B (1-308) were cloned in the pBIG1a biGBac vector using the biGbac cloning method (50). The kinase domain of Lck kinase (225-509) was synthesized (ThermoFisher Gene Art Gene Synthesis) in a PET151 vector backbone with an N-terminal cleavable Twin-strep and 6x his tag. It was then subcloned into a pLIB vector with a cleavable Twin-strep tag and 10x his tag. Final pLIB, pACEbac1 and pBIG1a constructs were transformed into DH10EmBacY cells (Geneva Biotech), with white colonies indicating successful generation of bacmids containing our genes of interest. Plasmid encoding TTC7B and FAM126A (1 to 308) was subcloned into a pOPT vector containing an N-terminal 10x histidine tag, followed by a 2x strep tag, followed by a tobacco etch virus (TEV) protease cleavage site (51). EFR3A (721 to 791) was subcloned into a pOPT vectors containing an N-terminal MBP tag in addition to the 10x histidine tag and 2x Strep tag. Nanobody F3IN was cloned into the pET22b vector backbone for inducible expression in the periplasm.GFP-SpyTag-(GS)_6_-EFR3A (721-791) was cloned into a pOPT vector with N-terminal 2x strep tag, 10x his tag and TEV cleavage site followed by the EGFP-Spytag with a 6 x GS linker before residues 721-791 of EFR3A (Referred to as EGFP-SpyTag-(GS)-EFR3A) . For expression in HEK293-AT1 cells, the F3IN nanobody sequence was amplified from the the pET22b-F3IN construct as template, using primers containing EcoRI and AgeI restriction sites. The fragment was then cloned into the iRFP-N1 plasmid using the same restriction digestion. The BRET constructs for measuring PI4KA PM recruitment and PM PI(4)P levels have been previously described (11, 52).

### Protein Expression

Plasmid containing the coding sequences for MBP-EFR3A (721-791) was expressed in BL21 DE3 C41 *Escherichia coli* and induced with 0.5 mM IPTG and grown at 37 °C for 4 h. The TTC7B-FAM126A (1-308), TTC7B-FAM126B (1-308) and chimeric TTC7B-EFR3A-FAM126A (1-308) constructs were expressed in BL21 DE3 C41 cells and induced with 0.1mM IPTG and grown at 21 °C for 20 hours. EGFP-SpyTag-(GS)-EFR3A was expressed in BL21 DE3 C41 cells induced with 0.1 mM IPTG and grown overnight at 18 °C. The F3IN construct was expressed in BL21 DE3 C41 cells and induced with 1mM IPTG and grown at 25°C for 20 hours. Cells were then harvested and centrifuged at 15,000*g*. Pellets were washed with PBS before being snap-frozen in liquid nitrogen, followed by storage at −80 °C.

Bacmid harboring TTC7A-FAM126A (1-308), TTC7A-FAM126B (1-308), PI4KA-TTC7B-FAM126A (1-308), PI4KA-TTC7A-FAM126A (1-308), ybbr-PI4KA-TTC7B-FAM126A (1-308), and LCK (225-509) were transfected into *Spodoptera frugiperda (Sf9) cells*, and viral stocks amplified for one generation to acquire a P2 generation final viral stock. Final viral stocks were added to *Sf9* cells in a 1:100 virus volume to cell volume ratio. Constructs were expressed for 65-72 hours before harvesting of the infected cells. Cell pellets were washed with PBS, flash frozen in liquid nitrogen, and stored at −80 °C.

### Protein Purification

#### F3IN purification

Nanobody F3IN was isolated from cell pellets by osmotic shock. Cell pellets were resuspended in 20 ml of TES buffer (0.2 M Tris pH 8.0, 0.5 mM ethylenediaminetetraacetic acid (EDTA), 0.5 M sucrose) with Protease Inhibitor Cocktail Set III and rotated for 45 min at 4°C. Twice the volume of TES/4 buffer was added to the cells and rotated for 45 min at 4°C, followed by centrifugation at 20,000*g* for 20 min at 4 C in a JA-20 rotor. The periplasmic extract was loaded onto a His-Trap HP column (Cytiva) equilibrated with Ni–NTA A buffer (20 mM Tris [pH 8.0], 100 mM NaCl, 20 mM imidazole [pH 8.0], 5% [v/v] glycerol, and 2 mM βME). The column was washed with 4 CV of high salt buffer (20 mM Tris [pH 8.0], 1 M NaCl, 5% [v/v] glycerol, and 2 mM bME), then 3-4 CV of Ni–NTA A buffer, followed by 3-4 CV of 6% Ni–NTA B buffer (20 mM Tris [pH 8.0], 100 mM NaCl, 200 mM imidazole [pH 8.0], 5% [v/v] glycerol, and 2 mM bME) before being eluted with 3-4 CV of 100% Ni–NTA B buffer. The protein elution was then buffer exchanged into PI4KA GFB (20 mM Imidazole [pH 7.0], 150 mM NaCl, 5% [v/v] glycerol and 0.5 mM Tris (2-carboxyethyl) phosphine (TCEP) and concentrated in a 10 kDa MWCO concentrator (Millipore Sigma). The concentrated protein was loaded onto the Superdex 75 Increase 10/300 GL (Cytiva) pre-equilibrated in GFB. Protein fractions were collected and concentrated in a 10 kDa MWCO concentrator (Millipore Sigma), flash frozen in liquid nitrogen, and stored at −80 °C.

#### MBP-EFR3A, EGFP-SpyTag-EFR3A purification

Cell pellets were lysed by sonication for 5 min in lysis buffer (20 mM Tris [pH 8.0], 100 mM NaCl, 5% [v/v] glycerol, 20 mM imidazole, 2 mM β-mercaptoethanol [bME]), and protease inhibitors [Millipore Protease Inhibitor Cocktail Set III, EDTA free]). Triton X-100 was added to 0.1% v/v, and the solution was centrifuged for 45 min at 20,000*g* at 1 °C (Beckman Coulter J2-21, JA-20 rotor). The supernatant was then loaded onto a 5 ml HisTrap column (Cytiva) that had been equilibrated in nickel–nitrilotriacetic acid (Ni–NTA) A buffer (20 mM Tris [pH 8.0], 100 mM NaCl, 20 mM imidazole [pH 8.0], 5% [v/v] glycerol, and 2 mM βME). The column was washed with 4 CV of high salt buffer (20 mM Tris [pH 8.0], 1 M NaCl, 5% [v/v] glycerol, and 2 mM βME), then 3-4 CV of Ni–NTA A buffer, followed by 3-4 CV of 6% Ni–NTA B buffer (20 mM Tris [pH 8.0], 100 mM NaCl, 200 mM imidazole [pH 8.0], 5% [v/v] glycerol, and 2 mM βME) before being eluted with 3-4 CV of 100% Ni–NTA B. The eluate was then loaded onto a 5 ml StrepTrapHP column (Cytiva) and washed with 3-4 CV of high salt gel filtration buffer (GFB) (20 mM HEPES [pH 7.5], 500 mM NaCl, 5% [v/v] glycerol and 1 mM Tris(2-carboxyethyl) phosphine [TCEP]), then 1 CV of GFB containing 2 mM ATP, 10 mM MgCl_2_, and 150 mM KCl, followed by 4 CV of GFB (20 mM HEPES [pH 7.5], 150 mM NaCl, 5% [v/v] glycerol and 1 mM Tris (2-carboxyethyl) phosphine [TCEP]). Protein was eluted with 3 CV of GFB containing 2.5 mM desthiobiotin and concentrated in a 10 kDa MWCO concentrator (Millipore Sigma). For the BLI competition experiments and TIRF microscopy (EGFP-SpyTag-EFR3A), the his-strep tag was cleaved using tobacco etch virus protease containing a stabilizing lipoyl domain (Lip-TEV). Concentrated protein was loaded onto the Superdex 75 Increase 10/300 GL (Cytiva) or the Superdex 200 Increase 10/300 GL (Cytiva) pre-equilibrated in GFB. Protein fractions were collected and concentrated in a 10 kDa MWCO concentrator (Millipore Sigma), flash frozen in liquid nitrogen, and stored at −80 °C.

#### TTC7-FAM126 (1-308) purification

Cell pellets were lysed by sonication for 5 min in lysis buffer (20 mM imidazole [pH 8.0], 100 mM NaCl, 5% [v/v] glycerol, 2 mM bME), and protease inhibitors [Millipore Protease Inhibitor Cocktail Set III, animal-free]. Triton X-100 was added to 0.1% v/v, and the solution was centrifuged for 45 min at 20,000*g* at 1 °C. The supernatant was then loaded onto a 5 ml HisTrap column (Cytiva) that had been equilibrated in nickel–nitrilotriacetic acid (Ni–NTA) A buffer (20 mM Tris [pH 8.0], 100 mM NaCl, 20 mM imidazole [pH 8.0], 5% [v/v] glycerol, and 2 mM βME). The column was washed with 3-4 CV Ni-NTA A buffer (20 mM imidazole [pH 8.0], 100 mM NaCl, 5% [v/v] glycerol, and 2 mM βME) followed by 3-4 CV of 6% Ni–NTA B buffer (450 mM Imidazole [pH 8.0], 100 mM NaCl, 5% [v/v] glycerol, and 2 mM βME) before being eluted with 3-4 CV of 100% Ni–NTA B. The eluate was then loaded onto a 5 ml StrepTrapHP column (Cytiva) and then washed with 3-4 CV of GFB (20 mM Imidazole [pH 7.0], 150 mM NaCl, 5% [v/v] glycerol and 0.5 mM Tris (2-carboxyethyl) phosphine (TCEP)). The His-strep tag was cleaved with Lip-TEV. TEV cleavage proceeded overnight following which the protein was eluted with GFB. Protein eluate was further concentrated in a 50 kDa MWCO concentrator (Millipore Sigma). Concentrated protein was loaded onto the Superdex 200 Increase 10/300 GL (Cytiva) or the Superose 6 Increase 10/300 GL (Cytiva) pre-equilibrated in GFB. Protein fractions from a single peak were collected and concentrated in 50 kDa MWCO concentrator (Millipore Sigma), flash frozen in liquid nitrogen and stored at -80 °C until further use.

#### PI4KA-TTC7A/B-FAM126A (1-308) purification

Sf9 pellets were resuspended in lysis buffer [20 mM imidazole pH 8.0, 100 mM NaCl, 5% glycerol, 2 mM βMe, protease (Protease Inhibitor Cocktail Set III, Sigma)] and lysed by sonication. Triton X-100 was added to 0.1% v/v final, and lysate was centrifuged for 45 min at 20,000*g* at 1 °C. The supernatant was then loaded onto a 5 ml HisTrap column (Cytiva) that had been equilibrated in nickel–nitrilotriacetic acid (Ni–NTA) A buffer (20 mM Tris [pH 8.0], 100 mM NaCl, 20 mM imidazole [pH 8.0], 5% [v/v] glycerol, and 2 mM bME). The column was washed with 3-4 CV of Ni-NTA A buffer (20 mM imidazole [pH 8.0], 100 mM NaCl, 5% [v/v] glycerol, and 2 mM βME) followed by 3-4 CV of 6% Ni–NTA B buffer (27 mM Imidazole [pH 8.0], 100 mM NaCl, 5% [v/v] glycerol, and 2 mM βME) before being eluted with 3-4 CV of 100% Ni–NTA B. Eluted protein was loaded onto a 5 ml StrepTrapHP column (Cytiva) pre-equilibrated GFB (20 mM imidazole pH 7.0, 150 mM NaCl, 5% glycerol [v/v], 0.5 mM TCEP). The His-strep tag was cleaved with Lip-TEV, and cleavage proceeded overnight. Cleaved protein was eluted with GFB and concentrated in a 50 kDa MWCO concentrator (MilliporeSigma). Concentrated protein was loaded onto the Superose 6 Increase 10/300 GL (Cytiva) pre-equilibrated in GFB. Protein fractions from a single peak were collected and concentrated in 50 kDa MWCO concentrator (Millipore Sigma), flash-frozen in liquid nitrogen and stored at -80 °C until further use.

#### LCK (225-509) purification

Cell pellets were lysed by sonication for 5 min in lysis buffer (20 mM Tris [pH 8.0], 100 mM NaCl, 5% [v/v] glycerol, 20 mM imidazole, 2 mM β-mercaptoethanol [bME]), and protease inhibitors [Millipore Protease Inhibitor Cocktail Set III, EDTA free]). Triton X-100 was added to 0.1% v/v final, and the solution was centrifuged for 45 min at 20,000*g* at 1 °C. The supernatant was then loaded onto a 5 ml HisTrap column (Cytiva) that had been equilibrated in nickel–nitrilotriacetic acid (Ni– NTA) A buffer (20 mM Tris [pH 8.0], 100 mM NaCl, 20 mM imidazole [pH 8.0], 5% [v/v] glycerol, and 2 mM bME). The column was washed with 4 CV of Ni–NTA A buffer, followed by 4 CV of 6% Ni–NTA B buffer (20 mM Tris [pH 8.0], 100 mM NaCl, 200 mM imidazole [pH 8.0], 5% [v/v] glycerol, and 2 mM bME) before being eluted with 4 CV of 100% Ni–NTA B. The eluate was then loaded onto a 5 ml StrepTrapHP column (Cytiva) and washed with 2.5 CV of GFB (20 mM HEPES [pH 7.5], 150 mM NaCl, and 0.5 mM Tris(2-carboxyethyl) phosphine [TCEP]. The His-strep tag was cleaved with Lip-TEV, and cleavage proceeded overnight. Cleaved protein was eluted with 2 CV of GFB and concentrated in a 10 kDa MWCO concentrator (MilliporeSigma). Concentrated protein was treated with 1 mM ATP and 2 mM MgCl_2_ for 1 minute at RT and then loaded onto the Superdex 75 Increase 10/300 GL (Cytiva) pre-equilibrated in GFB. Protein fractions were collected and concentrated in a 10 kDa MWCO concentrator (Millipore Sigma), flash frozen in liquid nitrogen, and stored at −80 °C until further use. The raw SDS-PAGE gel images for all purified proteins are shown in the source data.

#### SpyCatcher and EGFP-SpyTag

The his10-TEV-SUMO-KCK-SpyCatcher and his10-TEV-SUMO-KCK-EGFP-SpyTag(AHIVMVDAYKPTK*) fusion proteins were expressed in Rosetta (DE3) *E. coli*. Gene sequences used to recombinantly express SpyCatcher and SpyTag were inspired by previously described constructs (39). Bacteria were grown at 37°C in Terrific Broth to OD_600_ = 0.8. Cultures were then shifted to 18°C for 1 hour before inducing with 0.1 mM IPTG. Expression was allowed to continue for 18-21 hours before harvesting. Cells were lysed in 50 mM Na_2_HPO_4_ [pH 8.0], 400 mM NaCl, 10 mM imidazole, 4 mM BME, 1 mM PMSF, 100 μg/mL DNase by sonication. Lysate was centrifuged at 35000 *g* for 60 minutes in a Beckman JA-20 rotor at 4°C. NiNTA resin was added to the supernatant and allowed to bind for 2 hours, then washed with 50 mM Na_2_HPO_4_ [pH 8.0], 400 mM NaCl, 30 mM imidazole, 4 mM BME. Bound protein was eluted in 50 mM Na_2_HPO_4_ [pH 8.0], 400 mM NaCl, 500 mM imidazole, 4 mM BME. Peak fractions were pooled and loaded onto a desalting column equilibrated in 50 mM Na_2_HPO_4_ [pH 8.0], 400 mM NaCl, 20 mM imidazole, 5% glycerol, 1 mM BME. Peak fractions were pooled and incubated with SUMO protease (i.e. his6-SenP2) at 4°C overnight to cleave off the his10-TEV-SUMO tag. After 16-20 hours, the cleaved proteins were recirculated over NiNTA resin to capture the his6-SenP2 and the his10-TEV-SUMO tag. The flowthrough was concentrated in a 10 kDa MWCO Amicon-

Ultra concentrator before loading onto a 120 mL Superdex 75 column equilibrated in 20 mM Tris [pH 8.0], 200 mM NaCl, 10% glycerol, 1 mM TCEP. Peak fractions containing KCK-SpyCatcher were pooled and concentrated to 167 µM. The EGFP-SpyTag was loaded on a 24 mL Superdex 75 column equilibrated in 20 mM HEPES [pH 7.0], 150 mM NaCl, 10% glycerol, 1 mM TCEP. Peak fractions containing EGFP-SpyTag were pooled and concentrated to 484 µM. Both proteins were snap frozen in liquid nitrogen before storage at -80°C.

### Preparation of supported lipid bilayers

Supported lipid bilayers (SLBs) were created to maximize membrane fluidity at room temperature (i.e. 23°C). The following lipids were used to generate small unilamellar vesicles (SUVs) by hydration and extrusion: 1,2-dioleoyl-sn-glycero-3-phosphocholine (18:1 DOPC, Avanti #850375C), 1,2-dioleoyl-sn-glycero-3-phosphoethanolamine-N-[4-(p-maleimidomethyl)cyclohexane-carboxamide] (18:1 MCC-PE, Avanti #780201C), and phosphatidylinositol from bovine liver (PI, Avanti #840042C). To prepare the small unilamellar vesicles, a total of 2 µmoles lipids were combined in a 35 mL glass round bottom flask containing ∼2 mL of chloroform. Lipids were dried to a thin film using rotary evaporation with the glass round-bottom flask submerged in a 42°C water bath. After evaporating all the chloroform, the round bottom flask was place under a vacuum overnight at 23°C. The lipid film was then resuspended in 2 mL of 0.45 µm filtered 1x PBS pH 7.4 (Corning, cat# 46-013-CM), bring the lipid suspension to a final concentration of 1 mM. To generate 50 nm SUVs, the 1 mM total lipid mixtures was extruded through a 0.05 µm pore size 19 mm polycarbonate (PC) membrane (Cytiva Whatman, Cat# 800308) with filter supports (Cytiva Whatman, Cat# 230300) on both sides of the PC membrane. All lipid compositions used for supported lipid bilayer experiments contained the following molar percentages: 94% DOPC, 4% PI, 2% MCC-PE.

Supported lipid bilayers were formed on 25 x 75 mm coverglass (IBIDI, #10812) as previously described (53). In brief, coverglass was first cleaned with 2% Hellmanex III (ThermoFisher, Cat#14-385-864) and then etched with Piranha solution (1:3, hydrogen peroxide:sulfuric acid) for 15 minutes. Etched coverglass, washed extensively with MilliQ water again, is rapidly dried with nitrogen gas before adhering to a 6-well sticky-side chamber (IBIDI, Cat# 80608). SLBs were formed by flowing 0.25 mM total lipid concentration of 50 nm SUVs diluted in 1x PBS pH [7.4] into an assembled IBIDI chamber. SUVs were incubated in the IBIDI chamber for 30 minutes and then washed with 4 mL of PBS [pH 7.4] to remove non-absorbed SUVs. Membrane defects are blocked for 5 minutes with a solution of clarified 1 mg/mL beta casein (ThermoFisher, Cat# 37528) in 1x PBS [pH 7.4]. After blocking membranes with 1 mg/mL beta casein for 5 minutes, bilayers were washed with 4 mL of 1x PBS. Reaction/imaging buffer was exchanged into the chamber prior to each experiment.

Conjugation of SpyCatcher to supported membranes was achieved as previously described (54). Supported membranes containing 2% MCC-PE lipids were used to covalently couple SpyCatcher containing single N-terminal cysteine (i.e. thiol). For membrane coupling, 100 µL of 20 µM SpyCatcher was diluted in a 1x PBS [pH 7.4] and 0.1 mM TCEP buffer. This was incubated in an IBIDI chamber containing the SLB for 2 hours at 23°C. Unreacted MCC-PE lipids were then quench for 15 minutes 5 mM beta-mercaptoethanol (BME) diluted in 1x PBS [pH 7.4]. Quenched membranes were then washed and stored in 1x PBS [pH 7.4] until ready for imaging.

To attach SpyTag proteins, solutions containing 100 nM of either EGFP-SpyTag or EGFP-SpyTag-EFR3A were flowed into the chamber containing SpyCatcher membrane conjugated to MCC-PE lipids. The membrane surface density was monitored by TIRF microscopy to ensure an equivalent density of EGFP-SpyTag and EGFP-SpyTag-EFR3A became linked to the membrane. Based on fluorescence brightness a final density of 10-50 EGFP-SpyTag molecules per µm^2^ was achieved. After 20 minutes of SpyTag conjugation to SpyCatcher, unbound SpyTag proteins were flushed from the chamber with 1x PBS [pH 7.4]. These membranes were subsequently used to visualize EFR3A-mediated membrane recruitment of AF647-PI4KA in the absence and presence of nanobody, F3IN.

### Fluorescent labeling of ybbr-PI4KA-TTC7B-FAM126A

The DY647-coenzyme A (CoA) derivative was generated in-house by combining 15 mM DY647 C_2_ maleimide (Invitrogen, no. A20347) in dimethyl sulfoxide with 10 mM CoA (Sigma-Aldrich, no. C3144, MW = 785.33 g/mol) dissolved in 1× PBS to a final concentration of 7.5 mM DY647 maleimide and 5 mM CoA. This mixture was incubated overnight at 20°C in an amber tube. DY647-CoA conjugate was wrapped in parafilm and stored at −20°C. We labeled recombinant PI4KA-TTC7B-FAM126A (1-308) containing an N-terminal ybbR13 motif (DSLEFIASKLA) using Sfp transferase and DY647-CoA (55). Prior to the labelling of the protein, 10 μM DY647-CoA and 5 mM DTT in GFB (20 mM imidazole pH 7.0, 150 mM NaCl, 5% glycerol, 0.5 mM TCEP) were combined and incubated for 5 minutes at 20°C followed by 2 minutes on ice. Subsequently, chemical labeling was achieved by adding 5 μM ybbr-PI4KA-TTC7B-FAM126A (1–308), 4 μM Sfp-his6, and 10 mM MgCl_2_ and incubating the reaction overnight on ice. Excess DY647-CoA was removed using a HiTrap® desalting column preequilibrated in 3CV of GFB. The protein was then concentrated in a 50 kDa MWCO concentrator (Millipore Sigma) and loaded on a Superose 6 Increase 10/300 GL (Cytiva) equilibrated in GFB. Protein fractions were collected and concentrated in 50 kDa MWCO concentrator (Millipore Sigma), flash-frozen in liquid nitrogen and stored at -80 °C until further use.

### Visualization of AF647-PI4KA supported membrane localization dynamics by TIRF microscopy

To visualize AF647-PI4KA localization on supported lipid bilayers, the enzyme was diluted into buffer containing 20 mM HEPES [pH 7.0], 150 mM NaCl, 1 mM ATP, 5 mM MgCl_2_, 0.5 mM EGTA, 200 µg/mL beta casein, 20 mM BME, 20 mM glucose, 320 µg/mL glucose oxidase (Serva, #22780.01 *Aspergillus niger*), 50 µg/mL catalase (Sigma, #C40-100MG Bovine Liver), and 1 mM Trolox (Cayman Chemicals, cat #10011659). Trolox was prepared using a previously described protocol that utilized UV irradiation to drive the formation of a quinone species (56, 57). Membrane binding of AF647-PI4KA was visualized using an inverted Nikon Eclipse Ti2 microscope using a 100x Nikon (1.49 NA) oil immersion TIRF objective. Fluorescently labeled proteins were excited with either a 488 nm or 637 nm diode laser (OBIS laser diode, Coherent Inc. Santa Clara, CA) controlled with a Vortran laser drive with acousto-optic tunable filters (AOTF) control. The power output measured through the objective for single particle imaging was 1-2 mW. Excitation light was passed through the following quad pass dichroic filter cubes before illuminating the sample. Fluorescence emission was detected on the iXion Life 897 EMCCD camera (Andor Technology Ltd., UK) position after a Nikon emission filter wheel housing the following emission filters: ET525/50M and ET700/75M (Semrock). All in vitro biochemistry experiments were reconstituted at room temperature (23°C). Microscope hardware was controlled by Nikon NIS elements.

### Quantification of AF647-PI4KA membrane localization and binding frequency by smTIRF

To calculate the normalized membrane localization the fluorescence intensity of membrane bound AF647-PI4KA in the presence of membrane-tethered SpyCatcher and EGFP-SpyTag-EFR3A, at least 20 TIRF microscopy images from multiple fields of view were collected on an iXion Life 897 EMCCD camera. Before normalization, the background fluorescence intensity was subtracted to determine only the contribution of membrane bound AF647-PI4KA to the overall fluorescence signal. Intensities across multiple images were averaged to calculate the mean and standard deviation. The full data set was then normalized to average fluorescence intensity of membrane bound AF647-PI4KA measured in the absence of nanobody, F3IN.

To calculate the cumulative membrane binding frequency of AF647-PI4KA shown in Figure 8C, data was extracted from single molecule TIRF microscopy movies collected at 20 frames per second. Using the ImageJ/Fiji plugin, TrackMate, every AF647-PI4KA membrane docking event was time stamped as previously described (58). This allowed us to determine the exact frame when a AF647-PI4KA molecule was first detected, which was then scored as a single membrane binding event. Although every molecule AF647-PI4KA was tracked until membrane dissociation or photobleaching, only new AF647-PI4KA membrane docking events were counted to calculate the cumulative membrane binding frequency. All binding events for a single video were integrated over a 60 second period, which represents cumulative membrane binding measured in the presence of 1 nM AF647-PI4KA at equilibrium. Data shown in Figure 8C was collected with 3 technical replicates that were averaged. We set a threshold of 3 frames (i.e. 155 ms) as the minimum AF647-PI4KA dwell time to be considered an actual binding event. Molecules that docked on the membrane surface but remained immobile we filtered out. The threshold for being considered immobile was set for trajectories with a displacement ≤ 0.11 µm (1 pixel). This includes a fraction of AF647-PI4KA molecules that non-specifically absorbed to exposed glass or supported lipid bilayer defects.

### Phosphorylation analysis

For the phosphorylation of EFR3B, (20 μM) of protein was mixed with 1mM of ATP, 5mM of MgCl_2_, and 13 ug of LCK kinase. An unphosphorylated control containing equivalent amount of GFB (20 mM HEPES [pH 7.5], 150 mM NaCl, 5% [v/v] glycerol and 0.5 mM TCEP) instead of LCK kinase was also prepared. Reactions were incubated overnight on ice at 4°C. Subsequently, the protein mixtures were loaded onto a Superdex 200 10/300 GL (Cytiva) pre-equilibrated in GFB. Protein fractions from a single peak were collected and concentrated in 30 kDa MWCO concentrator (Millipore Sigma), flash-frozen in liquid nitrogen and stored at -80 °C until further use.

Phosphorylation of all proteins was confirmed using mass spectrometry and FragPipe v18.0 analysis. The LC-MS analysis of these samples was carried out using the same pipeline as used in the HDX-MS section below. The phosphorylated and non-phosphorylated peptide ratios were determined by generating extracted ion chromatograms (EIC) in the Thermo Xcalibur qual browser. The full analysis with EIC and MS data is provided in the source data.

### Lipid Vesicle Preparation

For inhibitor response assays of the PI4KA complexes, lipid vesicles containing 100% PI were prepared by mixing the lipid dissolved in organic solvent. The solvent was evaporated in a stream of argon following which the lipid film was desiccated in a vacuum for 1 hour. The lipids were resuspended in lipid buffer (20 mM HEPES pH 7.5, 100 mM KCl and 0.5 mM EDTA) and the solution was sonicated for 15 minutes. The vesicles were subjected to ten freeze thaw cycles and passed through an extruder (Avanti Research) with a 100 nm filter 21 times. The extruded vesicles were aliquoted and stored at −80°C.

### Lipid Kinase Assays

All lipid kinase activity assays employed the Transcreener ADP2 Fluorescence Intensity (FI) Assay (Bellbrook labs) which measures ADP production. For assays measuring inhibitor response, the final concentration of 100% PI vesicles was 0.25 mg/mL, ATP was 100 μM and inhibitors were 10 μM, 2 μM, 400 nM, 80 nM, 16 nM, or 3.2 nM (GSK-A1 only) (1% DMSO final). 2 μL of PI4KA complex solution at 2X final concentration (10 nM in 2X) was mixed with 2 μL substrate solution containing ATP, vesicles and inhibitors and the reaction was allowed to proceed for 60 min at 20°C. The reaction was stopped with 4 μL of 2X stop and detect solution containing Stop and Detect buffer, 8 nM ADP Alexa Fluor 594 Tracer and 93.7 μg/mL ADP2 Antibody IRDye QC-1 and incubated for 60 minutes. The fluorescence intensity was measured using an Agilent BioTek synergy H1 plate reader at excitation 585 nm and emission 626 nm. This data was normalised against a 0–100% ADP window made using conditions containing either 100 μM ATP or ADP with kinase buffer.

### Isolation of nanobodies from the yeast surface display library

The nanobodies were isolated from a yeast display library obtained from Dr. A.C. Kruse’s laboratory at Harvard Medical School and described in (33). The detailed protocol to isolate nanobodies from the yeast display library can be obtained from the Kruse laboratory website at: (https://kruse.hms.harvard.edu/sites/kruse.hms.harvard.edu/files/files/Nanobody_protoc <u>ol_0.pdf</u>)

The yeast cultures were grown in a selective dropout medium without tryptophan at 30°C. Nanobody expression was induced before each step by growing the cells in a dropout medium without tryptophan and with 2% galactose at 25°C for 48 to 72 hours. The library screening involved two rounds of magnetic-activated cell sorting (MACS) followed by two rounds of fluorescence-activated cell sorting (FACS) and a final round of MACS with a counter-selection step. The protein antigens (TTC7B-FAM126A (1-308) or chimeric TTC7B-EFR3A-FAM126A) were labelled using the Fluoreporter FITC protein labeling kit (Invitrogen F6434), the AlexaFluor-488 antibody labeling kit (Invitrogen A20181) or the Alexa Fluor 647 antibody labeling kit (Invitrogen A20186).

For MACS1, 1.1 x 10^10^ induced yeast cells were washed and re-suspended in 4.5ml selection buffer (20mM HEPES pH 7.5, 150mM sodium chloride, 0.1% (w/v) bovine serum albumin, 5mM maltose) and incubated with 500μl anti-FITC microbeads (Miltenyi 130-048-701) at 4°C for 40 minutes. After incubation with the magnetic beads, the cells were spun down and the excess unbound beads were removed with the supernatant. The cells were then resuspended in buffer and passed through a LD column (Miltenyi 130-042-901) to eliminate any yeast clones expressing nanobodies that bind non-specifically to the magnetic beads. The flow through from the LD column was spun down and stained in a solution of 1μM FITC-labelled TTC7B-FAM126A (1–308) in selection buffer for 1 hour at 4°C, spun down and re-suspended in 4.5ml selection buffer and 500μl anti-FITC microbeads, incubated for 20 minutes at 4°C and loaded on a LS column (Miltenyi 130-042-401). The cells binding to the column were eluted and grown for the next round of selection. The same protocol was used for the second round of MACS, except that we started with 2.5 x 10^9^ induced cells, the antigen was 38nM Alexa Fluor 647-labeled TTC7B-FAM126A (1-308) and anti-Alexa Fluor 647 microbeads were used (Miltenyi 130-091-395).

FACS selection was performed using a Cytoflex SRT cell sorter (Beckman Coulter). The induced cells were labeled with 1:200 AF647-labeled anti-HA antibody (Invitrogen 26183-A647) and 17nM FITC-labeled TTC7B-FAM126A (1-308) (FACS1) or 10nM AF488-labeled TTC7B-FAM126A (1-308) (FACS2) in selection buffer. Cells staining for both fluorophores were collected.

A final round of magnetic selection (MACS3) was performed after FACS2 to enrich for nanobodies that bind the TTC7B-FAM126A (1-308) at the EFR3A-binding interface. 1.5 x 10^8^ induced cells were stained with 20nM AlexaFluor647-labeled chimeric TTC7B-EFR3A-FAM126A and anti-AlexaFluor647 microbeads as described above and loaded on a LD column. The flow through was collected and stained with 20nM AlexaFluor647-labeled TTC7B-FAM126A (1-308) and anti-AlexaFluor647 microbeads and loaded on a LS column. The cells bound to the column were eluted and plated to isolate single colonies. The colonies were picked, grown and induced in a 96-well plate format for analysis by flow cytometry using a Cytoflex instrument (Beckman Coulter). 2 x 10^6^ induced cells from each colony were stained with 1:1000 AF488-anti-HA antibody to monitor nanobody expression levels and either 10nM AF647-labeled TTC7B-FAM126A (1-308) or 10nM AF647-labeled chimeric TTC7B-EFR3A-FAM126A. Clones that bound well to the TTC7B-FAM126A (1-308) but not the chimeric TTC7B-EFR3A-FAM126A were kept for subcloning and further analysis.

### Cryo-EM sample preparation and data collection

The PI4KA complex and F3IN were thawed and mixed at a 1:3 molar ratio and left to incubate on ice for 15 minutes. BS^3^ crosslinker was added with a final concentration of 0.25 mM and left to incubate on ice for 15 minutes followed by the addition of TRIS pH 7 for a final concentration of 25 mM to quench the reaction and incubated on ice for at least 15 minutes. C-Flat 2/1-3Cu-T-50 grids were glow-discharged for 25 s at 15 mA using a Pelco easiGlow glow discharger. 3 μL of the PI4KA complex and F3IN solution were applied to the grids at 0.75 mg/mL and vitrified using a Vitrobot Mark IV (Thermo Fisher) by blotting for 1.5 s at a blot force of −5 at 4°C and 100% humidity. Grids were screened at the UBC High-Resolution Macromolecular Cryo-EM (HRMEM) facility on a 200kV Glacios microscope (Thermo Fisher) equipped with a Falcon 3 camera (Thermo Fisher) to determine suitability for further data collection. Data was collected at the UBC High-Resolution Macromolecular Cryo-EM (HRMEM) facility on a 300kV Krios G2 microscope (Thermo Fisher) equipped with a Falcon 4i camera (Thermo Fisher) and Selectris energy filter (Thermo Fisher), using EPU software (Thermo Fisher). Data were collected at 165,000x magnification with a pixel size of 0.77 Å and a total dose of 50 e^−^/Å^2^, using a defocus range of −0.5 to −2.0 μm.

### Cryo-EM image processing

All data processing was carried out using cryoSPARC v4.5.2. Movies were imported with default settings. Patch motion correction output F-crop factor set to 1/2 was applied to all movies to align the frames. The contrast transfer function (CTF) of the resulting micrographs was estimated using the patch CTF estimation job with default settings. Final particle stack was manually curated to contain only micrographs with a CTF fit resolution less than or equal to 10.

To generate initial 3D models, 587,975 particles were picked from 4021 micrographs using a templates generated from a 3D reconstruction of the PI4KA complex from an independent dataset (without F3IN) lowpass filtered to 20 Å. Particles were extracted with a box size of 560 pixels and subjected to two rounds of 2D classification, removing duplicate particles, with 40 online-EM iterations, to remove junk and ice contamination. The resulting 69,526 particles underwent ab-initio reconstruction with two classes.

The complete dataset was then template picked as above (3,176,029 particles) and inspected with the inspect pics job, removing particles picking ice contamination. 2,262,222 particles were extracted with a box size of 560 pixels and subjected to two rounds of 2D classification with 40-online EM iterations. The remaining 657,836 particles underwent heterogenous refinement using the two ab-initio reconstructions from the small dataset. 329,699 particles from the class resembling the PI4KA complex underwent two rounds of homogeneous refinement followed by particle orientation rebalancing (rebalance percentile 80) prior to homogeneous refinement with C2 symmetry enforced. 282,982 particles then underwent C2 symmetry expansion followed by two rounds of masked local refinement with a mask covering the entire molecule. This generated a global reconstruction with an overall resolution of 3.54 Å based on the Fourier shell correlation (FSC) 0.143 criterion. To improve the quality of the TTC7B-F3IN interface, mask bases containing either TTC7B-F3IN were generated in ChimeraX and uploaded to cryoSPARC to generate masks used for masked local refinement and particle subtraction. The masked local refinements had their fulcrums centered at the TTC7B-F3IN interface, and generated refinements with an overall resolution of 3.66 Å (Local Refinement A) and 3.77 Å (Local Refinement B) based on the Fourier shell correlation (FSC) 0.143 criterion. A composite map was then generated using the global reconstruction and local refinement A and B in ChimeraX. The local resolutions of the three maps were determined with an FSC threshold of 0.143. The workflow to generate the final cryo-EM map is shown in **Fig S2**.

### HDX-MS sample preparation

HDX reactions examining TTC7B-FAM126A (1-308) in the presence and absence of F3IN were carried out in 10 µL reaction volumes containing 15 pmol of TTC7B-FAM126A (1.5 µM TTC7B-FAM126A) and 30 pmol of F3IN (3 µM F3IN in the bound state). The exchange reactions were initiated by the addition of 8.4 µL of D_2_O buffer (20 mM Imidazole pH 7, 150 mM NaCl) to 1.6 µL of protein (final D_2_O concentration of 78.9% [v/v]). Reactions proceeded for 3s, 30s, 300s, 3000s, and 10,000s at 18°C before being quenched with ice cold acidic quench buffer, resulting in a final concentration of 0.6M guanidine HCl and 0.9% formic acid.

HDX reactions examining the PI4KA complex in the presence and absence of F3IN were carried out in 10 µL reaction volumes containing 15 pmol of protein (1.5 µM PI4KA complex, 1.5 µM F3IN in the bound state). The exchange reactions were initiated by the addition of 7.5 µL of D_2_O buffer (20 mM Imidazole pH 7, 150 mM NaCl) to 2.5 µL of protein (final D_2_O concentration of 70.2 % [v/v]). Reactions proceeded for 3s, 30s, 300s and 3000s, at 18°C before being quenched with ice cold acidic quench buffer, resulting in a final concentration of 0.6M guanidine HCl and 0.9% formic acid post quench.

All conditions and timepoints were created and run in independent triplicate. All samples were flash frozen immediately after quenching and stored at -80°C.

### Protein digestion and MS/MS data collection

Protein samples were rapidly thawed and injected onto an integrated fluidics system containing a HDx-3 PAL liquid handling robot and climate-controlled (2°C) chromatography system (Trajan), a Dionex Ultimate 3000 UHPLC system, as well as an Exploris 120 Mass Spectrometer (ThermoFisher).The full details of the automated LC system are described in (59). The samples were run over an immobilized pepsin column (Affipro; AP-PC-001) at 200 µL/min for 4 minutes at 2°C. The resulting peptides were collected and desalted on a C18 trap column (Acquity UPLC BEH C18 1.7 µm column (2.1 × 5 mm); Waters 186004629). The trap was subsequently eluted in line with an ACQUITY 300Å, 1.7 μm particle, 100 × 2.1 mm BEH C18 UPLC column (Waters), using a gradient of 3-10% B (Buffer A 0.1% formic acid; Buffer B 100% acetonitrile) over 1.5 minutes, followed by a gradient of 10-25% B over 4.5 minutes, followed by a gradient of 25-35% B over 5 minutes, finally after 1 minute at 35% B a gradient of 35-80% B over 1 minute was used. Mass spectrometry experiments acquired over a mass range from *m/z* 150 to 2200 using an electrospray ionization source operated at a temperature of 200°C and a spray voltage of 4.5 kV. Same LC system, gradient and columns were used for all samples.

### Peptide identification

Peptides were identified from the non-deuterated samples using data-dependent acquisition following tandem MS/MS experiments (0.5 s precursor scan from *m/z* 150-2000; twelve 0.25 s fragment scans from *m/z* 150-2000 on QTOF system; precursor scan from *m/z* 300-1500, 4 data dependent scans from *m/z* 300-1500, dynamic exclusion set to n=1 with 10s exclusion time, 6s expected LC peak width on Orbitrap system). MS/MS datasets were analysed using FragPipe v18.0 and peptide identification was carried out by using a false discovery-based approach using a database of purified proteins and known contaminants (60, 61). MSFragger was used, and the precursor mass tolerance error was set to -20 to 20ppm. The fragment mass tolerance was set at 20ppm. Protein digestion was set as nonspecific, searching between lengths of 4 and 50 aa, with a mass range of 400 to 5000 Da.

### Mass Analysis of Peptide Centroids and Measurement of Deuterium Incorporation

HD-Examiner Software (Sierra Analytics) was used to automatically calculate the level of deuterium incorporation into each peptide. All peptides were manually inspected for correct charge state, correct retention time, appropriate selection of isotopic distribution, etc. Deuteration levels were calculated using the centroid of the experimental isotope clusters. Results are presented as relative levels of deuterium incorporation, and the only control for back exchange was the level of deuterium present in the buffer (78.9% or 70.2%). Differences in exchange in a peptide were considered significant if they met all three of the following criteria: ≥5% change in exchange, ≥0.4 Da difference in exchange, and a p value <0.01 using a two tailed student t-test. The entire HDX-MS dataset with all the values and statistics are provided in the source data. Samples were only compared within a single experiment and were never compared to experiments completed at a different time with a different final D_2_O level. The data analysis statistics for all HDX-MS experiments are provided in the source data according to the guidelines of (62). The HDX-MS proteomics data have been deposited to the ProteomeXchange Consortium via the PRIDE (63) partner repository with the dataset identifier PXD066213.

### Bio-layer interferometry (BLI)

The BLI measurements were conducted using a Fortebio (Sartorius) K2 Octet using fiber optic biosensors according to the protocols of (64). Anti–penta-His biosensors were loaded using purified F3IN, which had a 6x His tag on the C-terminus. The biosensor tips were preincubated in the BLI buffer (20 mM HEPES [pH 7.5], 150 mM NaCl, 0.01%, bovine serum albumin, and 0.002% Tween-20) for 10 min before experiments began. The sequence of steps in each assay was regeneration, custom, loading, baseline, association, and dissociation. Every experiment was done at 25 °C with shaking at 1000 rpm. Technical replicates were performed by using the same fiber tip and repeating the steps outlined previously. Regeneration was performed by exposing the tips to regeneration buffer (Glycine pH 1.5) for 5s and BLI buffer for 5s and repeating the exposure for 6 cycles. BLI buffer was used for the custom, baseline, and dissociation steps; these steps were performed in the same well for a given sample. To determine affinity constants, F3IN was diluted in BLI buffer to 100 nM and was loaded onto the anti penta-His biosensor tips. TTC7B-FAM126A or PI4KA complex were also diluted in BLI buffer and added to the appropriate association wells. Non-specific association was controlled by loading 100 nM of His-MBP onto the biosensor tips and subtracting its responses from the responses measured with F3IN. The data were fit to a 1:1 model using the “full (assoc and dissoc) setting”. Reported kinetic binding constants (K_D_, k_on_, and k_off)_ values were calculated by taking the geometric mean of binding constants calculated for individual conditions and replicates.

For the competition experiments, F3IN was diluted in BLI buffer to 100 nM and TTC7B-FAM126A or PI4KA complex was diluted in BLI buffer to 50 nM or 10 nM respectively. MBP-EFR3A was added to the association wells in increasing concentrations (0, 15, 30, 60, 120, 240 nM). All BLI experiments were performed using two technical replicates.

### Live-cell imaging using confocal microscopy

Cell culture - HEK293-AT1 cells were maintained in Dulbecco’s modified Eagle’s medium (DMEM—high glucose, sodium pyruvate) containing 10% FBS and 1% penicillin-streptomycin. The cell line was treated with Plasmocin prophylactic (InvivoGen, San Diego, CA) at 25 µg/ml for 1 week after thawing. The subsequent passages were maintained at 5 µg/ml of Plasmocin.

Live cell imaging - HEK293-AT1 cells (350,000 cells) were seeded on 30 mm glass bottom culture dishes (#1.5, Cellvis) pre-coated with 0.01% poly-L-lysine solution (Sigma). Cells were transfected next day with the indicated plasmid DNAs (GFP-PI4KA 50 ng/well and the EFR3A/TTC7B/FAM126A triple plasmid 100 ng/well) with or without 100 ng/well 3FIN-iRFP plasmid using Lipofectamine 2000 (0.5 μL/well; Invitrogen) following the manufacturer’s protocol. After 1 day of transfection, the media was replaced with 1 ml modified Krebs-Ringer buffer (containing 120 mM NaCl, 4.7 mM KCl, 2 mM CaCl_2_, 0.7 mM MgSO4, 10 mM glucose, and 10 mM Na-Hepes, adjusted to pH 7.4) and cells were observed at room temperature with a Zeiss confocal microscope (LSM 980) using a 63x Plan-Apochromat oil-immersion objective (N.A: 1.4).

### Live-Cell Measurements of PM PI(4)P levels and PI4KA PM recruitment using BRET based Biosensors

HEK293-AT1 cells (0.5x10^5^ cells/well) were seeded in a 200 μL total volume to white-bottom 96 well plates pre-coated with 0.01% poly-L-lysine solution (Sigma) and cultured overnight. Cells were then transfected with 150 ng of the PM-PI4KABRET biosensor (L10-mVenus-tPT2A-nLuc-PI4KA) using Lipofectamine 2000 (1 μL/well). Lipofection was done within Opti-MEM (40 μL/well) according to the manufacturer’s protocol, but with the slight modification of removing the media containing the Lipofectamine-complexed DNA and replacing it with complete culture medium at between 4-6 hrs post-transfection. Where indicated, 300 ng/well of either an empty vector (pcDNA3.1), or the 3FIN-iRFP plasmid was co-expressed with the EFR3B-P2A-TTC7B-T2A-FAM126A plasmid (11) together with the PM-PI4KABRET biosensor. For PI(4)P measurements, cells were transfected with 100 ng of the PM PI(4)P biosensor [L10-mVenus-tPT2A-nLuc-P4M(2x)] alone or in combination with the indicated amounts of either empty vector (pcDNA3,1) or the 3FIN-iRFP plasmid. BRET measurements were made at 37°C using a VANTAstar Multimode Microplate Reader (BMG LABTECH) with customized emission filters (540/40 nm and 475/20 nm). Between 20-24 hrs post-transfection, the cells were quickly washed before being incubated for 30 mins in 50 µL of modified Krebs-Ringer buffer (containing 120 mM NaCl, 4.7 mM KCl, 2 mM CaCl2, 0.7 mM MgSO4, 10 mM glucose, 10 mM HEPES, and adjusted to pH 7.4) at 37°C in an incubator (no CO_2_). After the pre-incubation period, the cell-permeable luciferase substrate, [NanoGlo, 40 µL, final concentration 5 µM, Promega for PI4KA BRET, or Coelenterazine H, 40 µL, final concentration 5 µM, Regis Technologies (Morton Grove, IL) for PI(4)P BRET], was added and the signal from the mVenus fluorescence and nLuc luminescence were recorded using 485 and 530 nm emission filters over a 45 min baseline BRET measurement (90 sec / cycle). Detection time was always 500 ms for each wavelength. BRET measurements are presented as the average of the stable basal BRET signals obtained after substrate addition during the entire measurement process of 45 minutes. All measurements were carried out in triplicate wells and repeated in three independent experiments. From each well, the BRET ratio was calculated by dividing the 530 nm and 485 nm intensities.

## Data availability

The EM data have been deposited in the EM data bank with accession numbers EMD-70826 (Composite map), EMD-70822 (Consensus map), EMD-70824 (Local Refinement A map), and EMD-70825 (Local Refinement B map), and the associated structural model has been deposited to the PDB with accession number (PDB: 9O6T) (Extended PDB ID: pdb_00009O6T). The MS proteomics data have been deposited to the ProteomeXchange Consortium via the PRIDE partner repository with the dataset identifier PXD066213 (63). All raw data in all figures are available in the source data excel file. All data needed to evaluate the conclusions in the paper are present in the paper and/or the supporting information.

## Supporting information

This article contains supporting information.

## Acknowledgements

Grids were prepared and data collected at the High Resolution Macromolecular Electron Microscopy (HRMEM) facility at the University of British Columbia (https://cryoem.med.ubc.ca). We thank Claire Atkinson, Amy Wo, Barathy Deivanayaga, Liam Worrall and Natalie Strynadka.

## Funding and additional information

J.E.B. is supported by the Canadian Institute of Health Research (PJT-195808). A.L.S. was supported by a CIHR CGS-D scholarship. S.D.H is supported by the National Science Foundation CAREER Award (MCB-2048060). The work of T.B and P.J was supported by the Intramural Research Program of the National Institutes of Health (NIH). The contributions of the NIH author(s) are considered works of the United States Government. The findings and conclusions presented in this paper are those of the author(s) and do not necessarily reflect the views of the NIH or the U.S. Department of Health and Human Services. Confocal imaging was performed in the Microscopy Core of NICHD with the kind assistance of Drs. Vincent Schram and Ling Yi. C.K.Y. is supported by CIHR (PJT-168907) and the Natural Sciences and Engineering Research Council of Canada (NGPIN-2024-04139). HRMEM is funded by the Canadian Foundation for Innovation and the British Columbia Knowledge Development Fund. J.A.C is supported by the Natural Sciences and Engineering Research Council of Canada (NGPIN-41822) and the Canadian Institute of Health Research (PJT-175120 AND PJT-175136).

## Conflict of interest

J.E.B. reports research contracts from Calico Life Sciences and OmniAB therapeutics. The authors declare that they have no conflicts of interest with the contents of this article.

## Notes

### Summary of Updates

Addition of cellular and TIRF microscopy data

## References

1. Balla, T. (2013) Phosphoinositides: Tiny lipids with giant impact on cell regulation. Physiological Reviews. 93, 1019–1137

2. Burke, J. E., Triscott, J., Emerling, B. M., and Hammond, G. R. V. (2023) Beyond PI3Ks: targeting phosphoinositide kinases in disease. Nat Rev Drug Discov. 22, 357–386

3. Hammond, G. R. V., and Burke, J. E. (2020) Novel roles of phosphoinositides in signaling, lipid transport, and disease. Current Opinion in Cell Biology. 63, 57–67

4. Batrouni, A. G., and Baskin, J. M. (2021) The chemistry and biology of phosphatidylinositol 4-phosphate at the plasma membrane. Bioorganic and Medicinal Chemistry. 40, 116190

5. de Rubio, R. G., Ransom, R. F., Malik, S., Yule, D. I., Anantharam, A., and Smrcka, A. V. (2018) Phosphatidylinositol 4-phosphate is a major source of GPCR-stimulated phosphoinositide production. Sci Signal. 11, eaan1210

6. Chung, J., Torta, F., Masai, K., Lucast, L., Czapla, H., Tanner, L. B., Narayanaswamy, P., Wenk, M. R., Nakatsu, F., and De Camilli, P. (2015) PI4P/phosphatidylserine countertransport at ORP5- and ORP8-mediated ER–plasma membrane contacts. Science. 349, 428–432

7. Lees, J. A., Zhang, Y., Oh, M. S., Schauder, C. M., Yu, X., Baskin, J. M., Dobbs, K., Notarangelo, L. D., De Camilli, P., Walz, T., and Reinisch, K. M. (2017) Architecture of the human PI4KIIIα lipid kinase complex. Proceedings of the National Academy of Sciences of the United States of America. 114, 13720–13725

8. Dornan, G. L., Dalwadi, U., Hamelin, D. J., Hoffmann, R. M., Yip, C. K., and Burke, J. E. (2018) Probing the Architecture, Dynamics, and Inhibition of the PI4KIIIα/TTC7/FAM126 Complex. Journal of Molecular Biology. 430, 3129–3142

9. Baskin, J. M., Wu, X., Christiano, R., Oh, M. S., Schauder, C. M., Gazzerro, E., Messa, M., Baldassari, S., Assereto, S., Biancheri, R., Zara, F., Minetti, C., Raimondi, A., Simons, M., Walther, T. C., Reinisch, K. M., and De Camilli, P. (2016) The leukodystrophy protein FAM126A (hyccin) regulates PtdIns(4)P synthesis at the plasma membrane. Nature Cell Biology. 18, 132–138

10. Nakatsu, F., Baskin, J. M., Chung, J., Tanner, L. B., Shui, G., Lee, S. Y., Pirruccello, M., Hao, M., Ingolia, N. T., Wenk, M. R., and De Camilli, P. (2012) Ptdins4P synthesis by PI4KIIIα at the plasma membrane and its impact on plasma membrane identity. Journal of Cell Biology. 199, 1003–1016

11. Suresh, S., Shaw, A. L., Pemberton, J. G., Scott, M. K., Harris, N. J., Parson, M. A. H., Jenkins, M. L., Rohilla, P., Alvarez-Prats, A., Balla, T., Yip, C. K., and Burke, J. E. (2024) Molecular basis for plasma membrane recruitment of PI4KA by EFR3. Science Advances. 10, eadp6660

12. Barlow-Busch, I., Shaw, A. L., and Burke, J. E. (2023) PI4KA and PIKfyve: Essential phosphoinositide signaling enzymes involved in myriad human diseases. Current Opinion in Cell Biology. 83, 102207

13. Suresh, S., and Burke, J. E. (2023) Structural basis for the conserved roles of PI4KA and its regulatory partners and their misregulation in disease. Advances in Biological Regulation. 90, 100996

14. Chen, R., Giliani, S., Lanzi, G., Mias, G. I., Lonardi, S., Dobbs, K., Manis, J., Im, H., Gallagher, J. E., Phanstiel, D. H., Euskirchen, G., Lacroute, P., Bettinger, K., Moratto, D., Weinacht, K., Montin, D., Gallo, E., Mangili, G., Porta, F., Notarangelo, L. D., Pedretti, S., Al-Herz, W., Alfahdli, W., Comeau, A. M., Traister, R. S., Pai, S.-Y., Carella, G., Facchetti, F., Nadeau, K. C., Snyder, M., and Notarangelo, L. D. (2013) Whole Exome Sequencing Identifies TTC7A Mutations for Combined Immunodeficiency with Intestinal Atresias. J Allergy Clin Immunol. 132, 656–664.e17

15. Agarwal, N. S., Northrop, L., Anyane-Yeboa, K., Aggarwal, V. S., Nagy, P. L., and Demirdag, Y. Y. (2014) Tetratricopeptide Repeat Domain 7A (TTC7A) Mutation in a Newborn with Multiple Intestinal Atresia and Combined Immunodeficiency. J Clin Immunol. 34, 607–610

16. Yang, W., Lee, P. p. w., Thong, M.-K., Ramanujam, T. M., Shanmugam, A., Koh, M.-T., Chan, K.-W., Ying, D., Wang, Y., Shen, J. j., Yang, J., and Lau, Y. l. (2015) Compound heterozygous mutations in TTC7A cause familial multiple intestinal atresias and severe combined immunodeficiency. Clinical Genetics. 88, 542–549

17. Jardine, S., Dhingani, N., and Muise, A. M. (2019) TTC7A: Steward of Intestinal Health. Cellular and Molecular Gastroenterology and Hepatology. 7, 555–570

18. Zara, F., Biancheri, R., Bruno, C., Bordo, L., Assereto, S., Gazzerro, E., Sotgia, F., Wang, X. B., Gianotti, S., Stringara, S., Pedemonte, M., Uziel, G., Rossi, A., Schenone, A., Tortori-Donati, P., van der Knaap, M. S., Lisanti, M. P., and Minetti, C. (2006) Deficiency of hyccin, a newly identified membrane protein, causes hypomyelination and congenital cataract. Nat Genet. 38, 1111–1113

19. Biancheri, R., Zara, F., Rossi, A., Mathot, M., Nassogne, M. C., Yalcinkaya, C., Erturk, O., Tuysuz, B., Di Rocco, M., Gazzerro, E., Bugiani, M., van Spaendonk, R., Sistermans, E. A., Minetti, C., van der Knaap, M. S., and Wolf, N. I. (2011) Hypomyelination and Congenital Cataract: Broadening the Clinical Phenotype. Archives of Neurology. 68, 1191–1194

20. Gazzerro, E., Baldassari, S., Giacomini, C., Musante, V., Fruscione, F., La Padula, V., Biancheri, R., Scarfì, S., Prada, V., Sotgia, F., Duncan, I. D., Zara, F., Werner, H. B., Lisanti, M. P., Nobbio, L., Corradi, A., and Minetti, C. (2012) Hyccin, the Molecule Mutated in the Leukodystrophy Hypomyelination and Congenital Cataract (HCC), Is a Neuronal Protein. PLoS ONE. 7, e32180

21. Berger, K. L., Cooper, J. D., Heaton, N. S., Yoon, R., Oakland, T. E., Jordan, T. X., Mateu, G., Grakoui, A., and Randall, G. (2009) Roles for endocytic trafficking and phosphatidylinositol 4-kinase III alpha in hepatitis C virus replication. Proc. Natl. Acad. Sci. U.S.A. 106, 7577–7582

22. Berger, K. L., Kelly, S. M., Jordan, T. X., Tartell, M. A., and Randall, G. (2011) Hepatitis C virus stimulates the phosphatidylinositol 4-kinase III alpha-dependent phosphatidylinositol 4- phosphate production that is essential for its replication. J. Virol. 85, 8870–8883

23. Adhikari, H., Kattan, W. E., Kumar, S., Zhou, P., Hancock, J. F., and Counter, C. M. (2021) Oncogenic KRAS is dependent upon an EFR3A-PI4KA signaling axis for potent tumorigenic activity. Nat Commun. 12, 5248

24. Kattan, W. E., Liu, J., Montufar-Solis, D., Liang, H., Brahmendra Barathi, B., van der Hoeven, R., Zhou, Y., and Hancock, J. F. (2021) Components of the phosphatidylserine endoplasmic reticulum to plasma membrane transport mechanism as targets for KRAS inhibition in pancreatic cancer. Proc Natl Acad Sci U S A. 118, e2114126118

25. Waring, M. J., Andrews, D. M., Faulder, P. F., Flemington, V., McKelvie, J. C., Maman, S., Preston, M., Raubo, P., Robb, G. R., Roberts, K., Rowlinson, R., Smith, J. M., Swarbrick, M. E., Treinies, I., Winter, J. J. G., and Wood, R. J. (2014) Potent, selective small molecule inhibitors of type III phosphatidylinositol-4-kinase α- but not β-inhibit the phosphatidylinositol signaling cascade and cancer cell proliferation. Chem. Commun. (Camb.). 50, 5388–5390

26. Bojjireddy, N., Botyanszki, J., Hammond, G., Creech, D., Peterson, R., Kemp, D. C., Snead, M., Brown, R., Morrison, A., Wilson, S., Harrison, S., Moore, C., and Balla, T. (2014) Pharmacological and genetic targeting of the PI4KA enzyme reveals its important role in maintaining plasma membrane phosphatidylinositol 4-phosphate and phosphatidylinositol 4,5-bisphosphate levels. J Biol Chem. 289, 6120–6132

27. Hamers-Casterman, C., Atarhouch, T., Muyldermans, S., Robinson, G., Hammers, C., Songa, E. B., Bendahman, N., and Hammers, R. (1993) Naturally occurring antibodies devoid of light chains. Nature. 363, 446–448

28. Jin, B.-K., Odongo, S., Radwanska, M., and Magez, S. (2023) NANOBODIES®: A Review of Diagnostic and Therapeutic Applications. Int J Mol Sci. 24, 5994

29. Wu, X., Chi, R. J., Baskin, J. M., Lucast, L., Burd, C. G., DeCamilli, P., and Reinisch, K. M. (2014) Structural Insights into Assembly and Regulation of the Plasma Membrane Phosphatidylinositol 4-Kinase Complex. Developmental Cell. 28, 19–29

30. Hornbeck, P. V., Zhang, B., Murray, B., Kornhauser, J. M., Latham, V., and Skrzypek, E. (2015) PhosphoSitePlus, 2014: mutations, PTMs and recalibrations. Nucleic Acids Res. 43, D512–20

31. Yaron-Barir, T. M., Joughin, B. A., Huntsman, E. M., Kerelsky, A., Cizin, D. M., Cohen, B. M., Regev, A., Song, J., Vasan, N., Lin, T.-Y., Orozco, J. M., Schoenherr, C., Sagum, C., Bedford, M. T., Wynn, R. M., Tso, S.-C., Chuang, D. T., Li, L., Li, S. S.-C., Creixell, P., Krismer, K., Takegami, M., Lee, H., Zhang, B., Lu, J., Cossentino, I., Landry, S. D., Uduman, M., Blenis, J., Elemento, O., Frame, M. C., Hornbeck, P. V., Cantley, L. C., Turk, B. E., Yaffe, M. B., and Johnson, J. L. (2024) The intrinsic substrate specificity of the human tyrosine kinome. Nature. 629, 1174–1181

32. Kwon, J., Kim, D. H., Park, J. M., Park, Y. H., Hwang, Y. H., Wu, H.-G., Shin, K. H., and Kim, I. A. (2016) Targeting Phosphatidylinositol 4-Kinase IIIα for Radiosensitization: A Potential Model of Drug Repositioning Using an Anti-Hepatitis C Viral Agent. Int J Radiat Oncol Biol Phys. 96, 867–876

33. McMahon, C., Baier, A. S., Pascolutti, R., Wegrecki, M., Zheng, S., Ong, J. X., Erlandson, S. C., Hilger, D., Rasmussen, S. G. F., Ring, A. M., Manglik, A., and Kruse, A. C. (2018) Yeast surface display platform for rapid discovery of conformationally selective nanobodies. Nat Struct Mol Biol. 25, 289–296

34. Shaw, A. L., Suresh, S., Parson, M. A. H., Harris, N. J., Jenkins, M. L., Yip, C. K., and Burke, J. E. (2024) Structure of calcineurin bound to PI4KA reveals dual interface in both PI4KA and FAM126A. Structure. 32, 1973–1983.e6

35. Abramson, J., Adler, J., Dunger, J., Evans, R., Green, T., Pritzel, A., Ronneberger, O., Willmore, L., Ballard, A. J., Bambrick, J., Bodenstein, S. W., Evans, D. A., Hung, C.-C., O’Neill, M., Reiman, D., Tunyasuvunakool, K., Wu, Z., Žemgulytė, A., Arvaniti, E., Beattie, C., Bertolli, O., Bridgland, A., Cherepanov, A., Congreve, M., Cowen-Rivers, A. I., Cowie, A., Figurnov, M., Fuchs, F. B., Gladman, H., Jain, R., Khan, Y. A., Low, C. M. R., Perlin, K., Potapenko, A., Savy, P., Singh, S., Stecula, A., Thillaisundaram, A., Tong, C., Yakneen, S., Zhong, E. D., Zielinski, M., Žídek, A., Bapst, V., Kohli, P., Jaderberg, M., Hassabis, D., and Jumper, J. M. (2024) Accurate structure prediction of biomolecular interactions with AlphaFold 3. Nature. 10.1038/s41586-024-07487-w

36. James, E. I., Murphree, T. A., Vorauer, C., Engen, J. R., and Guttman, M. (2022) Advances in Hydrogen/Deuterium Exchange Mass Spectrometry and the Pursuit of Challenging Biological Systems. Chem Rev. 122, 7562–7623

37. Masson, G. R., Jenkins, M. L., and Burke, J. E. (2017) An overview of hydrogen deuterium exchange mass spectrometry (HDX-MS) in drug discovery. Expert Opin Drug Discov. 12, 981–994

38. Skinner, J. J., Lim, W. K., Bédard, S., Black, B. E., and Englander, S. W. (2012) Protein hydrogen exchange: testing current models. Protein Sci. 21, 987–995

39. Zakeri, B., Fierer, J. O., Celik, E., Chittock, E. C., Schwarz-Linek, U., Moy, V. T., and Howarth, M. (2012) Peptide tag forming a rapid covalent bond to a protein, through engineering a bacterial adhesin. Proc Natl Acad Sci U S A. 109, E690–697

40. Hunyady, L., Baukal, A. J., Gaborik, Z., Olivares-Reyes, J. A., Bor, M., Szaszak, M., Lodge, R., Catt, K. J., and Balla, T. (2002) Differential PI 3-kinase dependence of early and late phases of recycling of the internalized AT1 angiotensin receptor. J Cell Biol. 157, 1211–1222

41. Kim, Y. J., Pemberton, J. G., Eisenreichova, A., Mandal, A., Koukalova, A., Rohilla, P., Sohn, M., Konradi, A. W., Tang, T. T., Boura, E., and Balla, T. (2024) Non-vesicular phosphatidylinositol transfer plays critical roles in defining organelle lipid composition. EMBO J. 43, 2035–2061

42. Bojjireddy, N., Guzman-Hernandez, M. L., Reinhard, N. R., Jovic, M., and Balla, T. (2015) EFR3s are palmitoylated plasma membrane proteins that control responsiveness to G-protein-coupled receptors. J Cell Sci. 128, 118–128

43. Koester, A. M., Geiser, A., Laidlaw, K. M. E., Morris, S., Cutiongco, M. F. A., Stirrat, L., Gadegaard, N., Boles, E., Black, H. L., Bryant, N. J., and Gould, G. W. (2022) EFR3 and phosphatidylinositol 4-kinase IIIα regulate insulin-stimulated glucose transport and GLUT4 dispersal in 3T3-L1 adipocytes. Bioscience Reports. 42, BSR20221181

44. Chung, J., Nakatsu, F., Baskin, J. M., and De Camilli, P. (2015) Plasticity of PI4KIIIα interactions at the plasma membrane. EMBO reports. 16, 312–320

45. Batrouni, A. G., Bag, N., Phan, H. T., Baird, B. A., and Baskin, J. M. (2022) A palmitoylation code controls PI4KIIIα complex formation and PI(4,5)P2 homeostasis at the plasma membrane. J Cell Sci. 135, jcs259365

46. Ulengin-Talkish, I., Parson, M. A. H., Jenkins, M. L., Roy, J., Shih, A. Z. L., St-Denis, N., Gulyas, G., Balla, T., Gingras, A.-C., Várnai, P., Conibear, E., Burke, J. E., and Cyert, M. S. (2021) Palmitoylation targets the calcineurin phosphatase to the phosphatidylinositol 4-kinase complex at the plasma membrane. Nat Commun. 12, 6064

47. Beghein, E., and Gettemans, J. (2017) Nanobody Technology: A Versatile Toolkit for Microscopic Imaging, Protein-Protein Interaction Analysis, and Protein Function Exploration. Front Immunol. 8, 771

48. Koenig, P.-A., Das, H., Liu, H., Kümmerer, B. M., Gohr, F. N., Jenster, L.-M., Schiffelers, L. D. J., Tesfamariam, Y. M., Uchima, M., Wuerth, J. D., Gatterdam, K., Ruetalo, N., Christensen, M. H., Fandrey, C. I., Normann, S., Tödtmann, J. M. P., Pritzl, S., Hanke, L., Boos, J., Yuan, M., Zhu, X., Schmid-Burgk, J. L., Kato, H., Schindler, M., Wilson, I. A., Geyer, M., Ludwig, K. U., Hällberg, B. M., Wu, N. C., and Schmidt, F. I. (2021) Structure-guided multivalent nanobodies block SARS-CoV-2 infection and suppress mutational escape. Science. 371, eabe6230

49. Fischer, B., Uchański, T., Sheryazdanova, A., Gonzalez, S., Volkov, A. N., Brosens, E., Zögg, T., Kalichuk, V., Ballet, S., Versées, W., Sablina, A. A., Pardon, E., Wohlkönig, A., and Steyaert, J. (2024) Allosteric nanobodies to study the interactions between SOS1 and RAS. Nat Commun. 15, 6214

50. Weissmann, F., Petzold, G., VanderLinden, R., Huis In’t Veld, P. J., Brown, N. G., Lampert, F., Westermann, S., Stark, H., Schulman, B. A., and Peters, J. M. (2016) BiGBac enables rapid gene assembly for the expression of large multisubunit protein complexes. Proceedings of the National Academy of Sciences of the United States of America. 113, E2564–E2569

51. Gibson, D. G., Young, L., Chuang, R. Y., Venter, J. C., Hutchison, C. A., and Smith, H. O. (2009) Enzymatic assembly of DNA molecules up to several hundred kilobases. Nature Methods. 6, 343–345

52. Tóth, J. T., Gulyás, G., Tóth, D. J., Balla, A., Hammond, G. R. V., Hunyady, L., Balla, T., and Várnai, P. (2016) BRET-monitoring of the dynamic changes of inositol lipid pools in living cells reveals a PKC-dependent PtdIns4P increase upon EGF and M3 receptor activation. Biochim Biophys Acta. 1861, 177–187

53. Hansen, S. D., Lee, A. A., Duewell, B. R., and Groves, J. T. (2022) Membrane-mediated dimerization potentiates PIP5K lipid kinase activity. Elife. 11, e73747

54. Duewell, B. R., Faris, K. A., and Hansen, S. D. (2024) Molecular basis of product recognition during PIP5K-mediated production of PI(4,5)P2 with positive feedback. J Biol Chem. 300, 107631

55. Yin, J., Lin, A. J., Golan, D. E., and Walsh, C. T. (2006) Site-specific protein labeling by Sfp phosphopantetheinyl transferase. Nat Protoc. 1, 280–285

56. Cordes, T., Vogelsang, J., and Tinnefeld, P. (2009) On the mechanism of Trolox as antiblinking and antibleaching reagent. J Am Chem Soc. 131, 5018–5019

57. Hansen, S. D., Huang, W. Y. C., Lee, Y. K., Bieling, P., Christensen, S. M., and Groves, J. T. (2019) Stochastic geometry sensing and polarization in a lipid kinase-phosphatase competitive reaction. Proc Natl Acad Sci U S A. 116, 15013–15022

58. Drew, E. E., Nyvall, H. G., Parson, M. A. H., Talus, R. K., Burke, J. E., and Hansen, S. D. (2025) SH2-mediated steric occlusion of the C2 domain regulates autoinhibition of SHIP1 inositol 5- phosphatase. J Biol Chem. 10.1016/j.jbc.2025.110788

59. Stariha, J. T. B., Hoffmann, R. M., Hamelin, D. J., and Burke, J. E. (2021) Probing Protein-Membrane Interactions and Dynamics Using Hydrogen-Deuterium Exchange Mass Spectrometry (HDX-MS). Methods Mol Biol. 2263, 465–485

60. Dobbs, J. M., Jenkins, M. L., and Burke, J. E. (2020) Escherichia coli and Sf9 Contaminant Databases to Increase Efficiency of Tandem Mass Spectrometry Peptide Identification in Structural Mass Spectrometry Experiments. J Am Soc Mass Spectrom. 31, 2202–2209

61. Kong, A. T., Leprevost, F. V., Avtonomov, D. M., Mellacheruvu, D., and Nesvizhskii, A. I. (2017) MSFragger: ultrafast and comprehensive peptide identification in mass spectrometry-based proteomics. Nat Methods. 14, 513–520

62. Masson, G. R., Burke, J. E., Ahn, N. G., Anand, G. S., Borchers, C., Brier, S., Bou-Assaf, G. M., Engen, J. R., Englander, S. W., Faber, J., Garlish, R., Griffin, P. R., Gross, M. L., Guttman, M., Hamuro, Y., Heck, A. J. R., Houde, D., Iacob, R. E., Jørgensen, T. J. D., Kaltashov, I. A., Klinman, J. P., Konermann, L., Man, P., Mayne, L., Pascal, B. D., Reichmann, D., Skehel, M., Snijder, J., Strutzenberg, T. S., Underbakke, E. S., Wagner, C., Wales, T. E., Walters, B. T., Weis, D. D., Wilson, D. J., Wintrode, P. L., Zhang, Z., Zheng, J., Schriemer, D. C., and Rand, K. D. (2019) Recommendations for performing, interpreting and reporting hydrogen deuterium exchange mass spectrometry (HDX-MS) experiments. Nat. Methods. 16, 595– 602

63. Perez-Riverol, Y., Bai, J., Bandla, C., García-Seisdedos, D., Hewapathirana, S., Kamatchinathan, S., Kundu, D. J., Prakash, A., Frericks-Zipper, A., Eisenacher, M., Walzer, M., Wang, S., Brazma, A., and Vizcaíno, J. A. (2022) The PRIDE database resources in 2022: a hub for mass spectrometry-based proteomics evidences. Nucleic Acids Res. 50, D543–D552

64. Bates, T. A., Gurmessa, S. K., Weinstein, J. B., Trank-Greene, M., Wrynla, X. H., Anastas, A., Anley, T. W., Hinchliff, A., Shinde, U., Burke, J. E., and Tafesse, F. G. (2025) Biolayer interferometry for measuring the kinetics of protein-protein interactions and nanobody binding. Nat Protoc. 20, 861–883

