## supplement for "Development of an inhibitory TTC7B selective nanobody that blocks EFR3 recruitment of PI4KA"

**This file includes:**

**Fig S1:** SEC traces of PI4KA-F3IN on a Superose 6 10/300 Increase column……….S1

**Fig S2:** Cryo-EM data processing…………………………………………………………S2

**Fig S3:** Raw BLI traces of EFR3B binding to TTC7A-FAM126A/B..…………………..S3

**Table S1:** Table S1. Cryo-EM data collection, refinement, and validation statistics….S4

**Table S2:** Binding constants of the different complexes………………………………...S5

**Table S3:** Key Reagents/Resources………………………………………………………S6

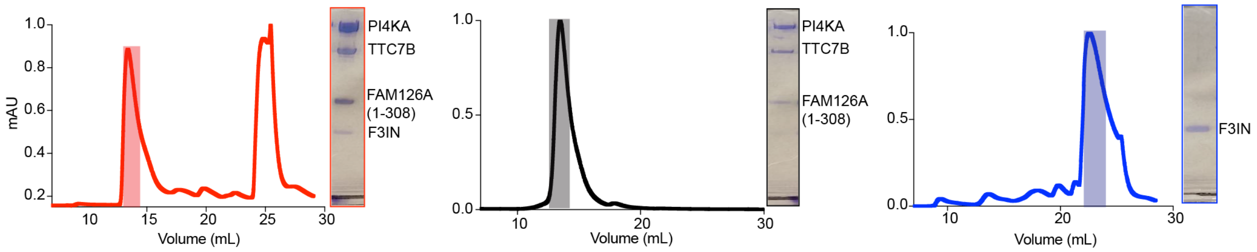

**Figure S1. SEC traces of PI4KA-F3IN on a Superose 6 10/300 Increase column.** (L) PI4KA-F3IN co-elution, (M) PI4KA alone, (R) F3IN alone

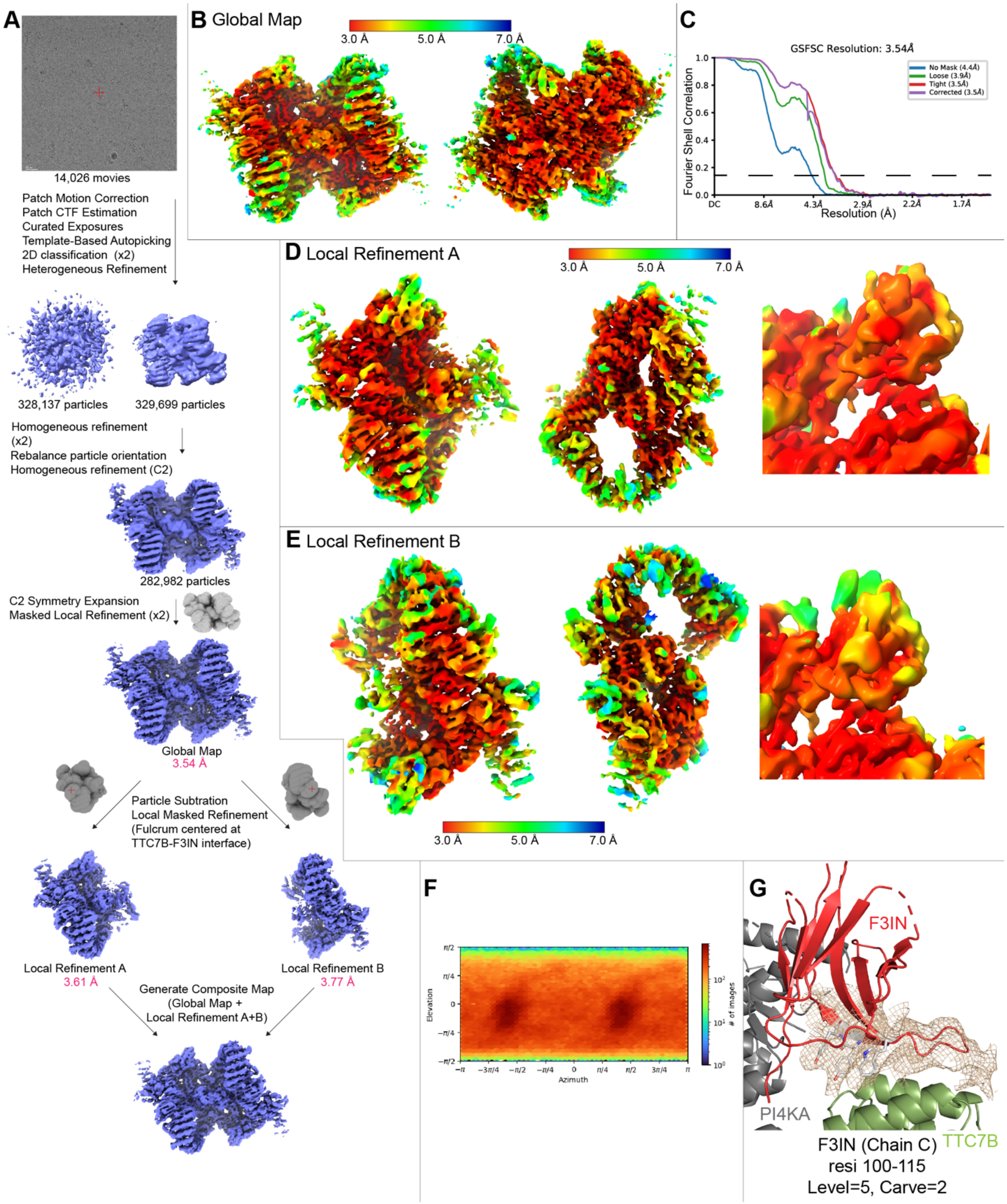

**Figure S2: Cryo-EM data processing**

(A) Cryo-EM data processing workflow showing a representative micrograph from screening on the 200 kV Glacios, and the processing strategy used to generate a 3D reconstruction of the PI4KA/TTC7B/FAM126A/F3IN complex.

(B) Global map coloured according to local resolution estimated using cryoSPARC v4.5.2 (FSC=0.143).

(C) Gold standard Fourier shell correlation coefficient (FSC) curve after auto tightening by cryoSPARC for the final map.

(D) Local refinement A map coloured according to local resolution estimated using cryoSPARC v4.5.2 (FSC=0.143).

(E) Local refinement B map coloured according to local resolution estimated using cryoSPARC v4.5.2 (FSC=0.143).

(F) Viewing direction distribution plot of particles in the global map output by cryoSPARC v4.5.2.

(G) Electron density of F3IN residues 100-115. Residues W101, Y112, and Y113 are shown.

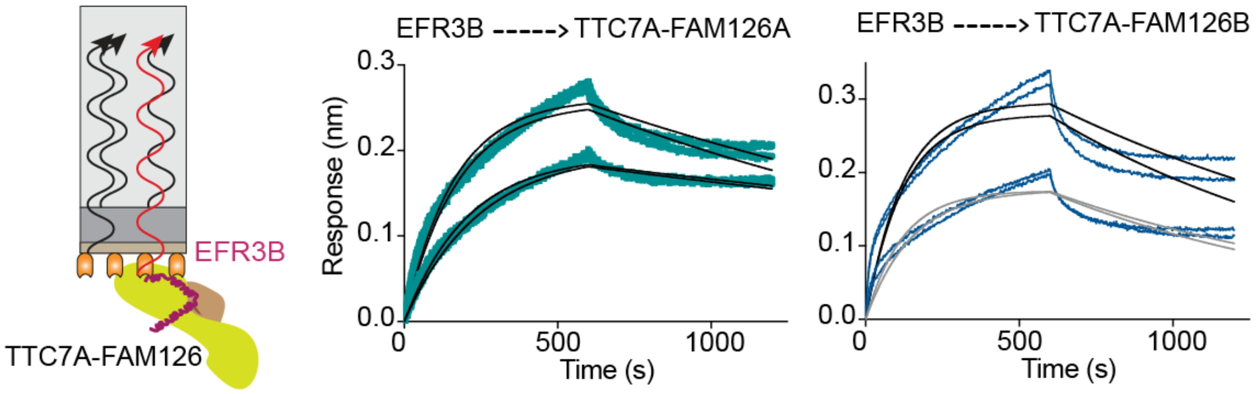

**Figure S3: Raw BLI traces of EFR3B binding to TTC7A-FAM126A/B.** (L) Cartoon of BLI experiment with EFR3B on the tip and TTC7A-FAM126A/B in solution. (C) Raw BLI traces of EFR3B binding with TTC7A-FAM126A (3.9 nM, 7.8 nM). (R) Raw BLI traces of EFR3B binding with TTC7A-FAM126B (15.6 nM, 20 nM).

**Table S1. Binding constants of the different complexes.**

| Immobilized ligand | Analyte | $K_D$ (M) | SD | $k_{on}$ (1/Ms) | SD | $k_{off}$ (1/s) | SD | Full $X^2$ | Sample size (n) |
|---|---|---|---|---|---|---|---|---|---|
| EFR3A | TTC7B-FAM126A | 4.08E-11 | 2.54E-11 | 4.10E+05 | 3.91E+04 | 1.68E-05 | 1.09E-05 | 0.250 | 4 |
| EFR3B | TTC7B-FAM126A | 4.05E-10 | 5.39E-11 | 5.40E+05 | 5.09E+04 | 2.19E-04 | 1.38E-05 | 0.190 | 4 |
| EFR3A | TTC7B-FAM126B | 2.98E-10 | 7.07E-11 | 4.07E+05 | 6.84E+04 | 1.21E-04 | 1.67E-05 | 0.427 | 4 |
| EFR3B | TTC7B-FAM126B | 3.45E-10 | 1.94E-10 | 5.95E+05 | 1.01E+05 | 2.05E-04 | 8.29E-05 | 0.248 | 4 |
| EFR3A | TTC7A-FAM126A | 7.18E-10 | 3.23E-10 | 3.59E+05 | 1.08E+05 | 2.58E-04 | 2.05E-04 | 0.300 | 4 |
| EFR3A | TTC7A-FAM126B | 4.73E-10 | 2.22E-10 | 9.28E+05 | 2.29E+05 | 4.39E-04 | 1.07E-04 | 0.257 | 4 |
| EFR3A | PI4KA-TTC7B-FAM126A | 4.31E-11 | 1.36E-11 | 8.55E+05 | 1.71E+05 | 3.69E-05 | 1.14E-05 | 0.124 | 3 |
| EFR3A | PI4KA-TTC7A-FAM126A | 3.39E-10 | 1.25E-10 | 8.06E+05 | 3.66E+05 | 2.73E-04 | 5.43E-05 | 0.138 | 3 |
| F3IN | TTC7B-FAM126A | 2.98E-09 | 8.26E-10 | 6.69E+05 | 2.76E+05 | 1.99E-03 | 3.84E-04 | 1.008 | 6 |
| F3IN | PI4KA-TTC7B-FAM126A | 2.97E-11 | 1.88E-11 | 3.79E+05 | 5.03E+04 | 1.12E-05 | 8.55E-06 | 1.345 | 5 |
| F3IN | PI4KA-TTC7A-FAM126A | 1.30E-09 | 1.39E-10 | 8.95E+05 | 9.49E+04 | 1.16E-03 | 2.47E-04 | 8.851 | 3 |

**Table S2. Cryo-EM data collection, refinement, and validation statistics**

| | PI4KA/TTC7B/FAM126A (1-308)/F3IN<br>EMD-70826<br>PDB 9O6T |
|---|---|
| **Data collection and processing** | |
| Magnification | 165,000 |
| Voltage (kV) | 300 |
| Electron exposure (e/ Å$^2$) | 50 |
| Defocus range ($\mu$M) | 0.5-2.0 |
| Pixel size (Å) | 0.77 |
| Symmetry imposed | C2 |
| Initial particle images (no.) | 3,176,029 |
| Final particle images (no.) | 282,982 |
| Map resolution (Å) | 3.54 |
| FSC threshold | 0.143 |
| Map resolution range (Å) | - |
| **Refinement** | |
| Initial model used (PDB) | 9BAX, AlphaFold3 |
| Model Resolution (Å) | 3.54 |
| FSC threshold | 0.143 |
| Map sharpening B factor | - |
| Model composition | |
| Non-hydrogen atoms | 42678 |
| Protein residues | 5344 |
| Ligands | 0 |
| *B*-factors | |
| Protein | 175.74 |
| Validation | |
| Mol probability score | 1.78 |
| Clashscore | 6.92 |
| Poor rotamers (%) | 1.59 |
| Ramachandran | |
| Favoured | 96.34 |
| Allowed | 3.62 |
| Outliers | 0.04 |
| R.M.S. deviations | |
| Bond lengths (Å) | 0.002 |
| Bond angles (°) | 0.489 |

### Table S3. Key Reagents/Resources

| REAGENT or RESOURCE | SOURCE | IDENTIFIER |
|---|---|---|
| **Bacterial and virus strains** | | |
| *E.coli* XL10-GOLD KanR Ultracompetent Cells | Agilent | 200317 |
| *E.coli* DH10EMBacY Competent Cells | Geneva Biotech | DH10EMBacY |
| C41(DE3) chemically competent cells | Lab stock | |
| **Chemicals, peptides, and recombinant proteins** | | |
| Deuterium Oxide 99.9% | Sigma Aldrich | 151882-10X1ML |
| $BS^3$ | Thermo Fischer Scientific | 21580 |
| ATP | Sigma | A7699-1g |
| $MgCl_2$ | Caledon Laboratory Chemicals | 4720-1 |
| Phosphatidylinositol (Liver PI) | Avanti Research | 840042C |
| GSK-A1 | SynKinase | CAS 1416334-69-4 |
| GSK-F1 | Adipogen | SYN-1220-M001 |
| Simeprevir | MedChemExpress | HY-10241 |
| **Critical Commercial assays** | | |
| Transcreener ADP2 FI Assay (1,000 Assay, 384 Well) | BellBrook Labs | 3013-1K |
| **Deposited Data** | | |
| Mass spectrometry proteomics data | https://www.ebi.ac.uk/pride/ | PXD066213 |
| PDB | https://www.rcsb.org/ | 9O6T (pdb_00009O6T) |
| EMDB | https://www.emdataresource.org/ | EMD-70826<br>EMD-70822<br>EMD-70824<br>EMD-70825 |
| **Recombinant DNA** | | |
| PI4KA-TTC7B-FAM126A (1-308) | (1) | GD177 |
| PI4KA-TTC7A-FAM126A (1-308) | This paper | SS171 |
| ybbr-PI4KA-TTC7B-FAM126A (1-308) | This paper | MJ345 |
| MBP-EFR3A (721-791) | (2) | MJ319 |
| MBP-EFR3A (721-791) | This paper | MJ326 |
| EGFP-SpyTag-(GS)-EFR3A 721-791 | This paper | SS187 |
| *his10-TEV-SUMO-KCK-EGFP-SpyTag* | | |
| his10-TEV-SUMO-KCK-SpyCatcher | | |
| MBP | (2) | AS42 |
| TTC7B- chimera EFR3A-FAM126A (1-308) | This paper | AS79 |
| TTC7B-FAM126A (1-308) WT | (1) | GD176 |
| TTC7B-FAM126A (1-308) WT – *E.coli* | (2) | AS29 |
| TTC7A-FAM126A (1-308) WT – *sf9* | This paper | SS145 |
| TTC7B-FAM126B (1-308) WT – *E.coli* | This paper | SS139 |
| TTC7A-FAM126B (1-308) | This paper | DA16 |
| Nanobody F3IN (protein sequence below) | This paper | SS170 (6C11) |

| | | |
|---|---|---|
| MKYLLPTAAAGLLLLAAQPAMAMAQVQLQESGGGLVQAGGSLRLSCAASGTIS ASDYMGWYRQAPGKERELVASIDGGGITNYADSVKGRFTISRDNAKNTVYLQM NSLKPEDTAVYYCAVDWILARYNFVIHYYWGQGTQVTVSSGSYPYDVPDYALE | | |
| LCK kinase (225-509) | This paper | MJ329 |
| EFR3Bha_gsgT2A_TTC7Bmyc_FlagFam126A | (2) | C3 |
| EGFP-PI4KA | (3) | |
| L10-mVenus-tPT2A-nLuc-PI4KA | (2) | PM-PI4KA$^{BRET}$ |
| F3IN-iRFP | This paper | F3IN-iRFP |
| **Software and algorithms** | | |
| cryoSPARC v4.5.2 | Structura Bio | https://cryosparc.com/ |
| Phenix-1.19.1 | Open source | https://phenix-online.org/ |
| COOT-0.9.4.1 | CCP4 | https://www2.mrc-lmb.cam.ac.uk/personal/pemsley/coot/ |
| PDBePISA | EMBL-EBI | https://www.ebi.ac.uk/pdbe/pisa/ |
| HDExaminer | Sierra Analytics | http://massspec.com/hdexaminer |
| Thermo Xcalibur Data Acquisition and Interpretation Software | Thermo Fisher Scientific | https://www.thermofisher.com/ |
| GraphPad Prism 7 | GraphPad | https://www.graphpad.com |
| FragPipe (v19.1) | Nesvizhskii Lab – University of Michigan | fragpipe.nesvilab.org |
| Adobe Illustrator 2021 | Adobe | https://www.adobe.com/products/illustrator.html |
| ESPript 3.0 | SBGrid consortium | https:espript.ibcp.fr |
| PyMOL | Schroedinger | http://pymol.org |
| ChimeraX | UCSF | https://www.cgl.ucsf.edu/chimerax/ |
| **Other** | | |
| C-flat Holey Thick Carbon Grid 2.0 μm hole 1.0 μm space 300 mesh | Electron Microscopy Studies | CFT312-100 |
| SF9 insect cells for expression | Expression Systems | 94-001S |
| Octet HIS1K Biosensors | Sartorius | 18-5120 |
| Enzymate Protein Pepsin Column, 300Å, 5μm, 2.1 mm X 30 mm | Waters | 186007233 |
| ProDx Pepsin Column F 10-32 | Trajan Scientific Americas | 359997870 |
| ACQUITY UPLC BEH C18 1.7 μm, 2.1 mm x 5 mm | Waters | 186004629 |
| ACQUITY UPLC Peptide BEH C18 Column, 300Å, 1.7 μm, 100 mm X 2.1 mm | Waters | 186003686 |

**Supplemental References**

1. Dornan, G. L., Dalwadi, U., Hamelin, D. J., Hoffmann, R. M., Yip, C. K., and Burke, J. E. (2018) Probing the Architecture, Dynamics, and Inhibition of the PI4KIIIα/TTC7/FAM126 Complex. *Journal of Molecular Biology*. **430**, 3129–3142
2. Suresh, S., Shaw, A. L., Pemberton, J. G., Scott, M. K., Harris, N. J., Parson, M. A. H., Jenkins, M. L., Rohilla, P., Alvarez-Prats, A., Balla, T., Yip, C. K., and Burke, J. E. (2024) Molecular basis for plasma membrane recruitment of PI4KA by EFR3. *Science Advances*. **10**, eadp6660
3. Hammond, G. R. V., Machner, M. P., and Balla, T. (2014) A novel probe for phosphatidylinositol 4-phosphate reveals multiple pools beyond the Golgi. *J Cell Biol*. **205**, 113–126
